# Glycosite Mapping and *in situ* Mass Spectrometry Imaging of MUC2 Glycopeptides via On-slide Digestion with Mucinase StcE

**DOI:** 10.1101/2024.09.16.613285

**Authors:** Sarah C. Lowery, Isabella P. Tran, Grace Grimsley, Rachel Stubler, Keira E. Mahoney, Taryn M. Lucas, Georgia Charkoftaki, Alvaro Santos-Neto, Nissi Varki, Vasilis Vasiliou, Richard R. Drake, Stacy A. Malaker

## Abstract

Many cancers are characterized by altered mucin expression and glycosylation, although the mechanistic relationship between tumor glycosylation and disease progression is not well-defined. Herein, our goal was to map specific mucin glycoforms in diseased tissue, enabling correlation of the tumor glycan profile with malignant features. To this end, we developed a workflow implementing on-tissue digestion with mucinase StcE, followed by matrix-assisted laser desorption ionization mass spectrometry imaging (MALDI-MSI) and liquid chromatography coupled to mass spectrometry (LC-MS). To optimize our workflow, we analyzed four different mucinous carcinomas derived from colon, esophageal, and salivary gland tissue.

Using this technique, we deduced the spatial distribution of StcE-generated O-glycopeptides within mucinous tumors using MALDI-IMS. Subsequent LC-MS analyses revealed the identity of different species detected in imaging experiments, in addition to comprehensively characterizing the mucinome and proteome of each tissue. Our coupled MS approach unveiled a striking mucin 2 (MUC2) expression pattern in two colorectal mucinous adenocarcinomas, in which different glycoforms clearly stratified regions within the tumor. Notably, our LC-MS experiments obtained near-complete sequence coverage over the mucin domains of MUC2, enabling glycoproteomic mapping of this canonical mucin in unprecedented depth. MUC2 glycosylation was dominated by the T and Tn antigens, with surprisingly little sialylation detected. However, O-glycans containing mono- and di-O-acetylated sialic acid were detected in low abundance. Finally, we obtained spectral evidence for an endogenous O-acetylated GalNAc, an O-glycan structure not previously reported in the literature. Overall, this proof-of-concept work underscores the potential of this technique to generate new research avenues in oncology and beyond.

**Significance Statement:** Aberrant mucin expression and glycosylation are hallmarks of cancer, but how these changes promote malignant processes are not well understood. Solid tumors are highly heterogeneous in their cellular and molecular composition, and many advanced spatial techniques have emerged in recent years to study the tumor microenvironment (TME) for better understanding disease progression. Spatially resolved glycoprotein analyses typically detect either the protein or glycan components, but not both. We developed a workflow using a dual mass spectrometry approach to map the location of intact glycopeptides in mucinous tumors, enabled by on-tissue digestion with the mucin-specific protease StcE. Future applications of this method on larger patient cohorts will enhance our understanding of glycans in malignancy, identify disease biomarkers, and define therapeutic targets.

## Introduction

Protein glycosylation is the most abundant post-translational modification (PTM), predicted to occur on over 50% of the human proteome.^1^ The majority of protein glycosylation can be classified as either N-linked or O-linked.^2^ N-linked glycosylation occurs primarily on asparagine residues within an NXS/T sequon, in which Asn is followed first by any amino acid except proline, and then a serine or threonine.^3^ All N-linked glycans are derived from the same chitobiose core, which is further extended by various glycosyltransferases in the endoplasmic reticulum.^4^ Although there are numerous types of O-linked glycosylation that occur on serine and threonine residues, most research has focused on intracellular O-GlcNAc or extracellular “mucin-type” O-GalNAc glycosylation. ^2,5^

The term “mucin-type” refers to the mucin family of glycoproteins, where this type of glycosylation is most abundant.^6^ Mucins, the primary protein component of mucus, are characterized by extremely dense O-GalNAc glycosylation in their proline-, threonine-, and serine-rich or “PTS” domains.^7^ The human proteome includes roughly 20 canonical mucins, which can be further classified as transmembrane or secreted.^8^ However, mucin domains are found in a diverse range of glycoproteins, including the T cell immunoglobulin and mucin-domain containing (TIM) family and platelet glycoprotein 1bα (GP1Bα).^9–11^ The unique properties of these domains are important for their varied functions in the formation of protective mucus barriers, regulation of proteolytic cleavage, and cell surface adhesion.^7,12,13^

Many cancers feature aberrant expression of different mucins, making them attractive diagnostic and therapeutic targets.^13^ For example, transmembrane mucin-16 (MUC16, also known as CA125) is overexpressed in most ovarian cancers and is an important serum biomarker for monitoring disease progression.^13^ Mechanistically, elevated expression of mucin motifs on the tumor surface have been shown to physically preclude immune cell engagement while simultaneously decreasing adhesion to surrounding tissue, facilitating metastatic invasion.^14–16^

The structure of glycans themselves can also play a significant role in pathological mechanisms, presenting an additional layer of complexity in studying mucins and other proteins within the glycocalyx.^17^ Increased expression of tumor-associated carbohydrate antigens (TACAs), including the sialylated Lewis-X (SLe^x^) and Lewis-A (SLe^a^) structures, is also observed on both N- and O-glycans.^18,19^ Specific O-glycan structures associated with cancer include the Tn, sialyl-Tn (STn), and Thomsen–Friedenreich (T) antigens.^19^ In fact, experimental tissue models have directly linked malignant phenotypes to the overexpression of truncated O-GalNAc glycans.^20^

However, the relationship between mucins and disease is not always well understood. For example, strong evidence exists for both protective and pathological functions of MUC2 in colorectal cancer (CRC).^21–23^ MUC2 is a secreted, gel-forming mucin that assembles into netlike assemblies that form a protective layer on intestinal epithelial cells.^24^ Histopathological analyses have shown diminished or even absent MUC2 expression in colorectal tumors compared to normal tissue,^22^ and MUC2 knockout mice lacking a protective intestinal mucus layer spontaneously develop colon tumors.^23^ On the other hand, mucinous carcinomas of the colon often overexpress MUC2.^25^ Mucinous colorectal adenocarcinoma, a distinct CRC subtype, is characterized by mucus content that exceeds 50% of the total tumor volume.^26^ Approximately 10-15% of all diagnosed CRC falls into this subcategory,^27^ which can be highly invasive and often responds poorly to traditional therapeutics.^26,28^ Mucinous carcinomas can also form in other epithelial tissues such as esophagus, pancreas, breast, and ovary, among others;^25,29^ the prognostic significance of mucinous differentiation appears to differ by tumor tissue origin.^30^

The exact molecular mechanisms governing the development and progression of mucinous tumors are poorly understood.^26^ Although glycosylation is essential for their structure and function, no canonical mucin has been sequenced at the glycoproteomic level to date. In fact, few studies have attempted to characterize both the expression and O-glycan profile of mucins in cancerous tissue, much less define the specific sites of glycosylation on their protein backbones. Liquid chromatography coupled to mass spectrometry (LC-MS) is the technique of choice for most glycoproteomic analyses, as it allows for comprehensive, untargeted detection of proteins in complex samples and can be used to localize glycans to specific amino acids.^31,32^ While LC-MS can comprehensively identify the various glycoforms present in a homogenized sample, such “bulk” analyses lose biologically relevant spatial information. Visualizing specific mucin glycoforms in malignant tissue could therefore improve our understanding of tumor physiology and inform therapeutic design.

In recent years, the high spatial heterogeneity within the tumor microenvironment and its relevance to disease progression has been documented extensively.^33^ Many techniques have been employed to understand the spatial distribution of mucins in various diseases, though all are accompanied by a few drawbacks. Histological dyes represent a simple, relatively inexpensive approach for mucin visualization in tissue. Alcian blue (AB)-periodic acid-Schiff (PAS) staining detects acidic and neutral mucins, respectively, although polysaccharides like glycogen are also stained.^34^ Catalytically inactivated mucinases, which are proteases selective for mucins, can be conjugated to reporter molecules for mucin-selective staining, but do not reveal the identities of the underlying mucins.^35^ More detailed analyses of specific mucin proteins and glycan structures typically requires antibody-based techniques such as immunohistochemistry (IHC).^36,37^ Recent advances in multiplexed antibody-based imaging, including technologies such as co-detection by indexing (CODEX)^38^ and multiplexed ion beam imaging by time of flight (MIBI-TOF)^39^ have allowed for simultaneous detection of multiple proteins (currently up to roughly 50) with high sensitivity and resolution. Although powerful, such approaches rely on prior knowledge of targets and the availability of well-characterized antibodies against them.

Matrix-assisted laser-desorption mass spectrometry imaging (MALDI-MSI) is a technique that is commonly employed to analyze glycans within preserved biological tissues with high spatial resolution.^40^ This technique is routinely used to map different analytes, including metabolites, lipids, and glycans.^41^ The latter is restricted to N-glycans, owing to the availability of the universal N-glycosidase, PNGaseF. To date, no universal O-glycosidase has been characterized, limiting spatial analyses of O-glycans to targeted methods such as lectin staining and IHC.

Mucinases, as described above, are a class of proteolytic enzymes specific for the unique bottlebrush-like structure of mucin domains. We hypothesized here that on-tissue mucinase digestion followed by MALDI-MSI could allow for visualization of mucin glycopeptide distribution. However, in MALDI-MSI, analytes are typically identified only by their intact precursor mass. This works well for species like N-glycans, whose modular structures limit them to a specific set of masses.^40,41^ In contrast, accurate identification of mucinase-generated O-glycopeptides with MS is impossible without tandem mass spectrometry. Given that we previously demonstrated the correlation between high-resolution MALDI imaging of N-glycans and LC-MS analysis of glycopeptides extracted from a serial tissue section in canine glioma,^42^ we reasoned that a similar approach could be employed here.

Our proof-of-concept experiments characterized colon, esophageal, and salivary gland carcinomas in this manner. Most of the detected O-glycopeptides were modified by known cancer antigens and many were found exclusively in benign or tumor tissue sections. Some of these tumor-associated O-glycopeptides were diffusely distributed within the carcinomas, while others localized to well-defined clusters. This work demonstrated that mucinase-derived O-glycopeptides determined by coupled MALDI-MSI and intact glycoproteomics can provide spatial distribution of dysregulated species that stratify distinct tumor regions. Furthermore, our approach enabled glycan mapping of tumor-associated MUC2 for the first time, and we achieved nearly complete sequence coverage across its two mucin domains. Overall, we present this workflow to identify mucin O-glycopeptides as potential diagnostic and/or therapeutic targets for mucinous carcinomas.

## Results & Discussion

### MALDI-IMS Can Map Glycopeptides Generated by On-tissue StcE Digestion

We analyzed four mucinous carcinomas with our coupled-MS approach: two from the colon, one from the esophagus, and one from the salivary gland. The colon tumors were resected from the same patient, while the esophageal and salivary gland tissue were collected from two different individuals. Both of the mucinous colorectal adenocarcinomas were classified as Stage II due to infiltration of neighboring muscle tissue.^43^ Additionally, the tumor regions bordered non-mucinous malignant tissue, and benign colon was present in each sample (**Figures S1 & S2**). Colon 1 also contained visually prominent adipose, vasculature, and lymphoid tissue (**Figure S1**). The mucinous esophageal tumor was adjacent to the junction of the columnar and squamous epithelia (**Figure S3**), a hotspot for the development of carcinomas.^44^ This sample may be a metastasis rather than a primary tumor, as the malignant tissue appeared to infiltrate from below and did not connect to the surface epithelium (**Figure S3**). Similarly, the salivary gland carcinoma might not represent a primary tumor. Salivary gland tumors are rarely mucinous,^45,46^ and the malignant region in this sample resided in the lymph node adjacent to parotid tissue, together supporting the characterization of this sample as a metastasis (**Figure S4**).

Using these tissues, we developed a MALDI-MSI workflow to image StcE-generated glycopeptides (**Figure 1A**). Briefly, tissue slides were de-waxed and subjected to low-pH antigen-retrieval, after which PNGaseF was applied using an automatic sprayer. Slides were rinsed to remove released N-glycans prior to on-slide StcE digestion, followed by matrix application and MALDI-MSI data acquisition. For each tissue, imaging experiments revealed numerous species that stratified tumor from non-tumor tissue (**Figures 1B-E**). Some of the masses detected in MALDI-MSI experiments corresponded to free N-glycans, despite rinsing slides after PNGaseF treatment, thus indicating a need for additional workflow optimization. Nearly all ions detected in high abundance outside of the tumor boundaries were N-glycans, although we identified multiple N-glycan species within the boundaries of the salivary gland tumor (**Figure S5**).

**Figure 1.**
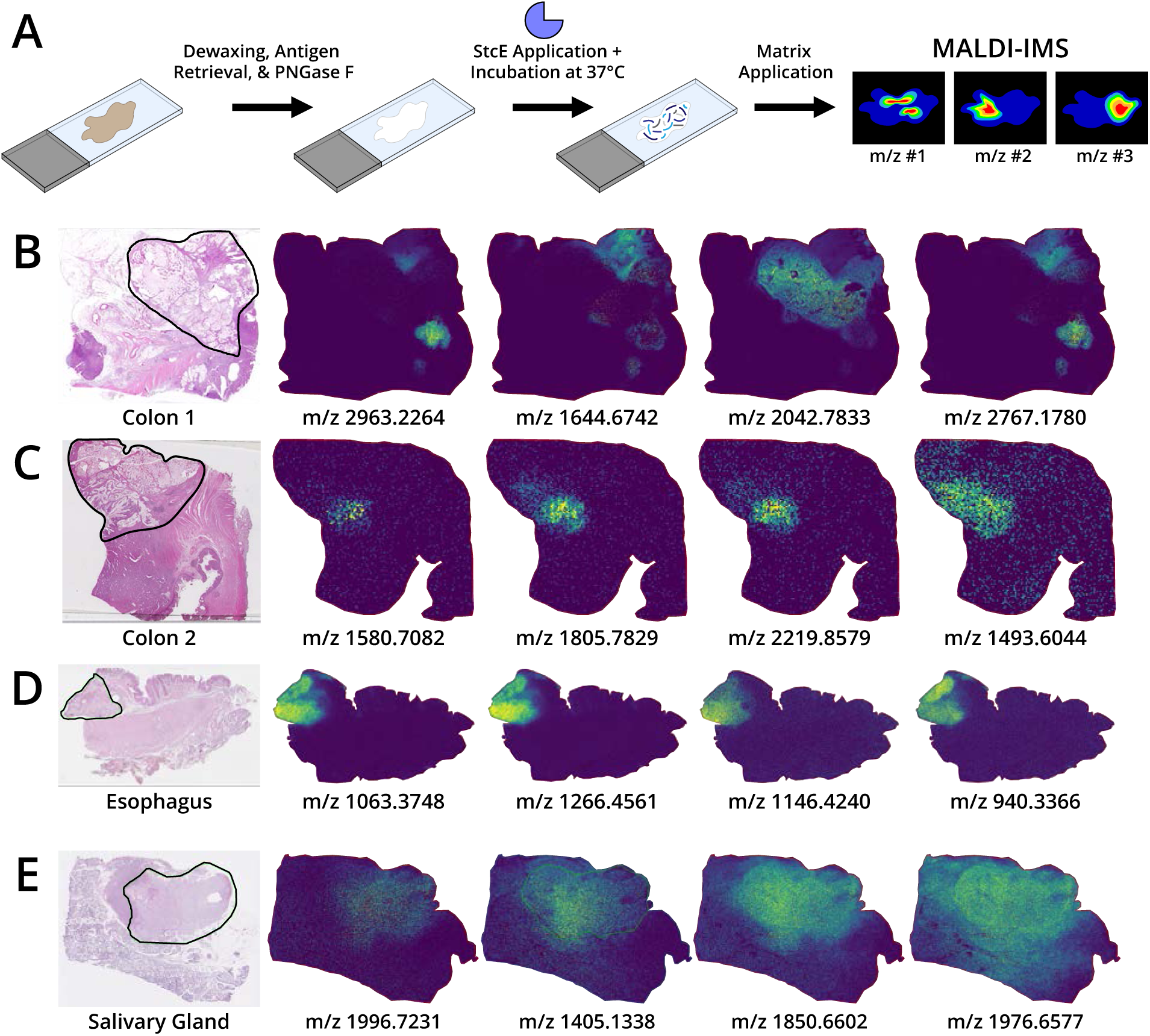
MALDI-IMS of tumor sections treated with PNGaseF and mucinase StcE. (A) FFPE tissue sections were dewaxed, antigen retrieved, and treated with PNGaseF. Following multiple rinses to remove released N-glycans, slides were subjected to on-slide digestion with StcE. After application of CHCA matrix, StcE-derived O-glycopeptides were spatially analyzed with MALDI-IMS. Tumor regions outlined on H&E-stained tissues and MALDI-IMS species representing distribution patterns observed (B) Colon 1, (C) Colon 2, (D) Esophagus, and (E) Salivary Gland samples.

Notably, different tumor-localized ions displayed distinct expression patterns even within the diseased tissue (**Figures 1B-E**). In Colon 1 and Colon 2, certain species localized to dense clusters, while others appeared more evenly distributed across the tumor region (**Figures 1B & 1C**). Ions from the esophageal tumor were generally detected across the entire tumor region, although some were more abundant along the edge next to the smooth muscle (**Figure 1D**). Most of the species detected in the salivary gland sample were diffusely distributed across the tumor, with some spillover into the adjacent parotid tissue (**Figure 1E**).

### Optimized LC-MS Sample Preparation Facilitates Tumor [Glyco]proteome Characterization

All slides used for LC-MS experiments were prepared as for MALDI-IMS prior to the matrix deposition step. The tumor and non-tumor regions were then outlined using a hydrophobic barrier pen (**Figure S6**). This allowed for a rough comparison in the (glyco)proteome between the malignant and adjacent tissues. Initially, we treated slides with StcE, followed by on-slide trypsin digestion, after which we extracted glycopeptides and unmodified peptides using a series of aqueous and organic solvents. After de-salting, we performed a hydrophilic interaction chromatography (HILIC)-based glycopeptide enrichment using polyhydroxyethyl aspartamide (PHEA) solid phase (**Materials & Methods**). However, search algorithms performed particularly poorly when analyzing enriched glycopeptide data, likely due to the enormous search space needed to accommodate the entire human proteome, non-specific proteolytic cleavage, multiple possible glycosites, and numerous glycan structures. Many of the glycopeptide backbone sequences identified in these searches were incorrect, requiring manual sequencing, a time-consuming approach that is untenable for large datasets.

Thus, we extracted the StcE-derived glycopeptides from each tissue to reduce sample complexity. Following a de-salting step, we acquired glycoproteomic data via LC-MS. When intact N- and O-glycopeptides are fragmented by collision-based techniques, the glycan’s labile glycosidic bonds are broken and its functional groups are further dissociated from the hexose ring, producing characteristic oxonium ions in the low-mass region. These fragments are collectively referred to as the HexNAc “fingerprint,” which is always present in glycopeptide MS2 spectra following collision-based dissociation.^31^ Unfortunately, we observed very weak glycopeptide signal, indicated by the intensity of the HexNAc diagnostic ion at *m/z* 204.0867 across the LC-MS gradient (**Figure S7a-c**). Searching this data also produced suboptimal results, with the low number of identifications likely stemming from weak MS2 spectra.

We determined that further saturating the tissue surface with StcE prior to extraction improved glycopeptide recovery and detection in downstream LC-MS based on the stronger *m/z* 204.0867 trace in these samples (**Figure S7a-c**). The HexNAc signal was particularly intense in LC-MS analyses tumor-derived glycopeptide samples, although lower levels were detected for samples collected from the non-tumor regions (**Figure S8a-b**). We compared glycopeptide recovery using this “double-StcE” and StcE-trypsin-PHEA workflows, the latter of which was used in our first experiment. An additional StcE digestion was not performed for the StcE-trypsin-PHEA workflow to prevent LC-MS signal saturation with tryptic peptides from StcE itself. Although the HexNAc traces using each method had similar profiles, ion intensity was stronger in the double-StcE glycopeptide sample (**Figure S9a-c**). This difference might stem from minor sample loss during PHEA enrichment or from ion suppression of glycopeptides by more readily ionized tryptic peptides during LC-MS acquisition. Thus, our optimized LC-MS sample preparation workflow opted for a second on-slide StcE treatment and extraction of glycopeptides prior to trypsin digestion.

After collecting glycopeptides in the double-StcE workflow, we dried the slides completely before depositing trypsin and digesting for 3-4 hours (**Figure 2a**). We then extracted and enriched the unmodified peptides from each tissue. By using the same PHEA enrichment method as in early experiments, we hoped to enrich non-mucinous O-glycopeptides from tryptic digests, but we observed only negligible HexNAc signal in these runs. Future work should test whether on-slide digestion with an O-glycoprotease, like the *Pseudomonas aeruginosa* immunomodulating protease (IMPa),^47^ enhances O-glycoproteomic coverage of non-mucin glycoproteins.

**Figure 2.**
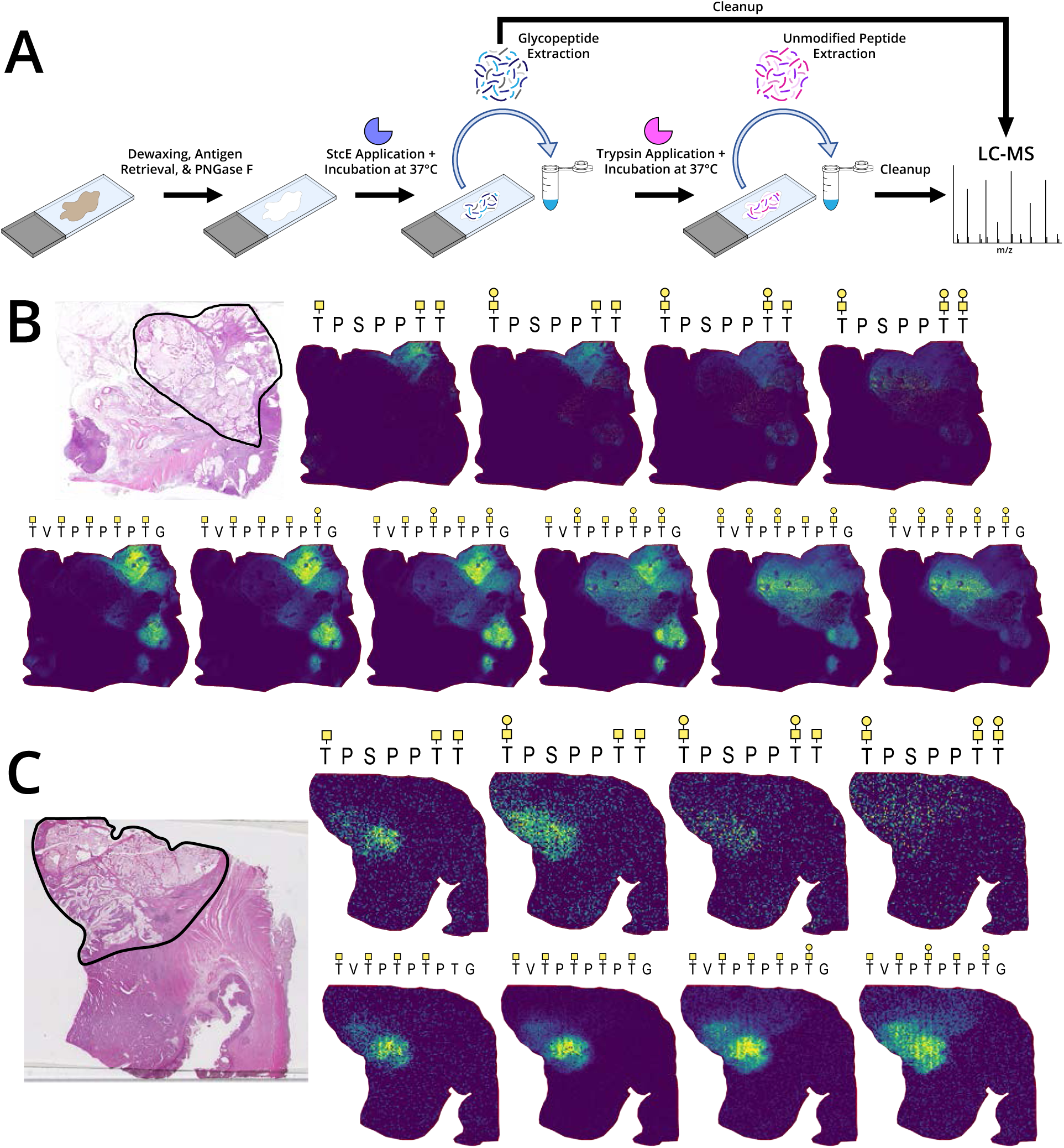
Identification of StcE-derived O-glycopeptides via LC-MS. (A) Adjacent tissue sections were dewaxed, antigen-retrieved, and PNGaseF-treated. Following rinse steps and on-slide digestion with sprayer-applied StcE, the tissue surface was saturated with pipette-applied StcE to maximize O-glycopeptide generation. After O-glycopeptide extraction, an on-slide trypsin digestion was performed, and unmodified peptides were collected and fractioned with a HILIC-based approach. All peptide samples were then analyzed with LC-MS. (B) The expression profile of MUC2 glycoforms in Colon 1 varies based on relative amounts of the T and Tn antigens, with the high levels of the former displaying a more diffuse pattern and the latter more clustered to specific areas. The sequence TPSPPTT is from the first PTS domain in MUC2, and TVTPTPTPTG is from the second. (C) MUC2 glycoforms in Colon 2 exhibit a similar spatial pattern as in Colon 1.

Subsequent analyses of tryptic peptide fractions collected during PHEA enrichment revealed the comprehensive proteome within each tissue (**Dataset S1**), allowing us to curate focused protein databases used to analyze glycopeptide data. This tailored database approach greatly enhanced the sensitivity, accuracy, and speed of glycoproteomic search algorithms by drastically reducing the size of the search space. We manually inspected MS spectra to validate all software-identified glycopeptides in the tumor LC-MS data (**Dataset S2**). First, we confirmed the peptide backbone sequence and total glycan composition using higher energy collision dissociation (HCD) MS2 spectra. We then localized glycans to specific Ser and Thr residues by assessing fragments in MS2 spectra from electron transfer dissociation (ETD) or hybrid electron transfer/higher-energy collision dissociation (EThcD). When search algorithm results were incorrect, the correct glycopeptide was determined by manual peptide sequencing. Following confident identification in one sample, glycopeptide MS spectra were extracted from the raw LC-MS data of the other tissues and similarly evaluated. This approach allowed for the most thorough and accurate glycoproteomic comparison between tumors, accounting for minor differences in spectral quality for each sample.

We detected non-glycosylated tryptic peptides for numerous canonical mucins from which no glycopeptides were identified (**Dataset S1**). MUC1 overexpression is estimated to occur in over half of all cancers,^13^ and we detected this transmembrane mucin in both of the colorectal cancers. Notably, we did not identify any glycopeptides from MUC1. StcE cleaves mucins at so-called “T*XT” motifs, in which a glycosylated Ser or Thr is followed by any amino acid (X) and then another Ser or Thr.^48^ Cleavage occurs C-terminally to the “X” residue, and StcE can accommodate glycosylation on the second Ser or Thr in this sequence. Although MUC1 contains very few putative StcE cleavage sites in its tandem repeats, numerous StcE-sensitive sequences can be found outside this region. Another transmembrane mucin, MUC12, was identified using unmodified tryptic peptides in both colorectal tumors (**Dataset S1**). While expression of MUC12 is normally high in the gastrointestinal tract, particularly in the colon, its downregulation is associated with colorectal cancer and ulcerative colitis,^49,50^ which could help explain why we were only able to validate a single glycopeptide from this transmembrane mucin (**Dataset S2**).

In the esophageal tumor, we identified tryptic peptides from MUC5AC, the major secretory mucin of the respiratory tract and upper digestive system. We also identified a single peptide from MUC5B, another secreted airway mucin. The backbone sequences of both MUC5AC and MUC5B contain numerous Ser and Thr within TXT motifs, so the lack of MUC5AC and MUC5B glycopeptides in our data was surprising. Like MUC2, these secreted mucins assemble into complex, higher-order structures that may render the protein backbone inaccessible to StcE.^51^ MUC18, also known as melanoma cell adhesion molecule (MCAM) or CD146, was identified in all samples. This transmembrane mucin is relatively small and contains few putative StcE sites, potentially explaining the lack of glycopeptides in our results.

Using the unmodified peptide data, we identified 7 proteins that were exclusively identified in the tumor regions of each tissue (**Table S1**). Previous studies have linked aberrant expression of many of these proteins to malignant processes. For example, upregulation of acid ceramidase (Q13510) has been observed in a variety of different cancer types, and in some instances, drug-induced downregulation of this protein was shown to enhance the efficacy of chemotherapy *in vitro* and in mouse models.^52^ Another protein in this category, latent-transforming growth factor beta-binding protein 2 (LTBP2, Q14767), interacts with various ECM proteins in elastic fiber-rich tissues,^53^ and its expression was previously linked to worse patient outcomes in head and neck squamous cell carcinoma.^54^

Other ECM-related proteins also displayed notable expression patterns, particularly those related to collagen assembly and organization. Collagen remodeling is a well-documented feature of many solid tumors, often resulting in increased tissue stiffness and protection from immune cells.^55,56^ Although most collagens were consistently identified across tissues, the alpha-1 chain of type X collagen (Q03692) was detected in the salivary and both colon tumor regions (**Dataset S1**). Overexpression of type X collagen is observed in a variety of cancers and is has been associated with poor patient prognosis.^57–59^ Unlike most other mammalian glycoproteins, collagens can be glycosylated on hydroxylysine (Hyl) residues.^5^ Hyl are typically decorated with unextended or mono-glucosylated galactoses thought to play critical roles in collagen stability and fibril formation.^60^ The colon tumors contained procollagen-lysine, 2-oxoglutarate 5-dioxygenase 1 (PLOD1), and procollagen galactosyltransferase 1 (GT251), which catalyze the hydroxylation and subsequent galactosylation of lysine residues in collagens.^60^ Both steps can also be performed by a single enzyme, such as the multifunctional procollagen lysine hydroxylase and glycosyltransferase LH3 (PLOD3),^60^ which we detected in the tumor regions of Colon 2 and the esophagus. These data suggest that changes in collagen glycosylation may be pathologically significant in mucinous carcinomas, although rigorous follow-up is necessary to confirm this relationship.

### Integration of MALDI-MSI and LC-MS Data Enables Visualization of MUC2 Glycoforms

We matched MALDI-MSI heat maps to LC-MS glycopeptide identifications by manually matching ions between raw data files. Because the MALDI and electrospray ionization (ESI) mechanisms tend to produce different species of precursor ions,^61^ all *m/z* values detected in LC-MS analyses were first converted to *m/z* values likely to be found in MALDI-MSI data, or vice versa. MALDI tends to produce singly-charged precursor ions derived from adduction with a sodium ion, potassium ion, or proton.^62,63^ Ultimately, all species identified in our IMS experiments were singly charged and adducted to one or two sodium ions. On the other hand, ESI of mucinase-derived O-glycopeptides primarily produced protonated precursor ions that were doubly- or triply-charged (**Dataset S2**).

The sensitivity of LC-MS experiments superseded that of MALDI-MSI, as evidenced by the high number of identified O-glycopeptides lacking MALDI heat maps. This is partially due to the acquisition parameters used in MALDI-MSI experiments, including variations in the scan range. The maximum *m/z* values used in our experiments were 4000 (Colon 1), 2500 (Colon 2), and 3000 (Esophagus & Salivary Gland). A narrower MS scan range can enhance detection of ions within the *m/z* limits, but a more restricted *m/z* window will likely reduce the number of species identified. For example, the singly charged, mono-sodiated precursor mass exceeded the upper scan limit of 2500 *m/z* for 68 out of 575 O-glycopeptides identified in Colon 2 (**Dataset S2**). The disparity in MS sensitivity can also be attributed to differences in instrumentation. Intact glycopeptides suffer from poor ionization efficiency,^32^ an effect that might be more pronounced with MALDI compared to ESI. Further optimization of matrix composition and other components of sample preparation might improve glycopeptide ionization and detection by MALDI-MSI.

The vast majority of glycopeptides identified in the colorectal and esophageal tumors were derived from MUC2 (**Dataset S2**). Our coupled-MS approach revealed visually striking expression patterns of different MUC2 glycoforms in the colorectal tissues (**Figure 2B**). In Colon 1, Tn-only MUC2 glycopeptides generally clustered to two regions within the tumor, adjacent to fibrotic or non-mucinous malignant colon tissue. Tn-only glycoforms of TPSPPTT were also detected adjacent to lymphoid tissue and another area of non-mucinous malignant colon (**Figures 2B, S1, S10, & S11**). MUC2 glycoforms decorated with both T and Tn antigens were more diffusely distributed across the mucinous tumor area, and T-only glycoforms localized closer to the tumor edge opposite that of the Tn-only clusters, adjacent to adipose tissue (**Figure 2B, S1, S10, & S11**). We observed a similar expression pattern in Colon 2, with Tn-only glycopeptides concentrated at a single site bordering non-mucinous adenocarcinoma tissue and T-containing species spread over more of the tumor region (**Figure 2C, S2, S10, & S12**). Notably, StcE-derived O-glycopeptides were only identified within tumor boundaries, suggesting mucinous O-glycopeptides can define the borders of mucinous carcinomas with high specificity. For both colorectal tissues, we detected tumor-associated species with MALDI-MSI that could not be identified with LC-MS data, although the total number of spectra in this category was relatively low (**Figure S13**).

### StcE Facilitates in-depth O-Glycosite Mapping of Tumor-associated MUC2

To calculate MUC2 sequence coverage, we compiled all backbone sequences from identified glycopeptides as well as any unmodified peptides from tryptic digests. Additionally, because MUC2 results were nearly identical between Colon 1 and Colon 2, we consolidated results from these samples. We obtained 54.1% sequence coverage across the full MUC2 sequence, and 12% coverage outside of the mucin domains. The limited proteomic coverage outside of the mucin domains likely stemmed from the high number of disulfide-bonded cysteines combined with the lack of reduction and alkylation steps in our workflow. However, we obtained 91.2% coverage over its two PTS domains (**Figure 3A**), a particularly exciting result as no canonical mucin has been characterized at the glycoproteomic level to date. Most of the MUC2 glycopeptides identified in the colorectal tumors were decorated with a combination of T and Tn antigens, although we detected evidence indicating the presence of core 2, fucosylated, and sialylated O-glycans that could not be localized to specific Ser or Thr residues (**Dataset S2, Figure S14a**). In the first, shorter PTS domain, we confidently localized only Tn and T antigen structures, with the exception of one core 2 structure mapped to a single site (**Figure 3B**). Glycan structures localized on peptide sequences from the second PTS domain were significantly more diverse (**Figure 3C**).

**Figure 3.**
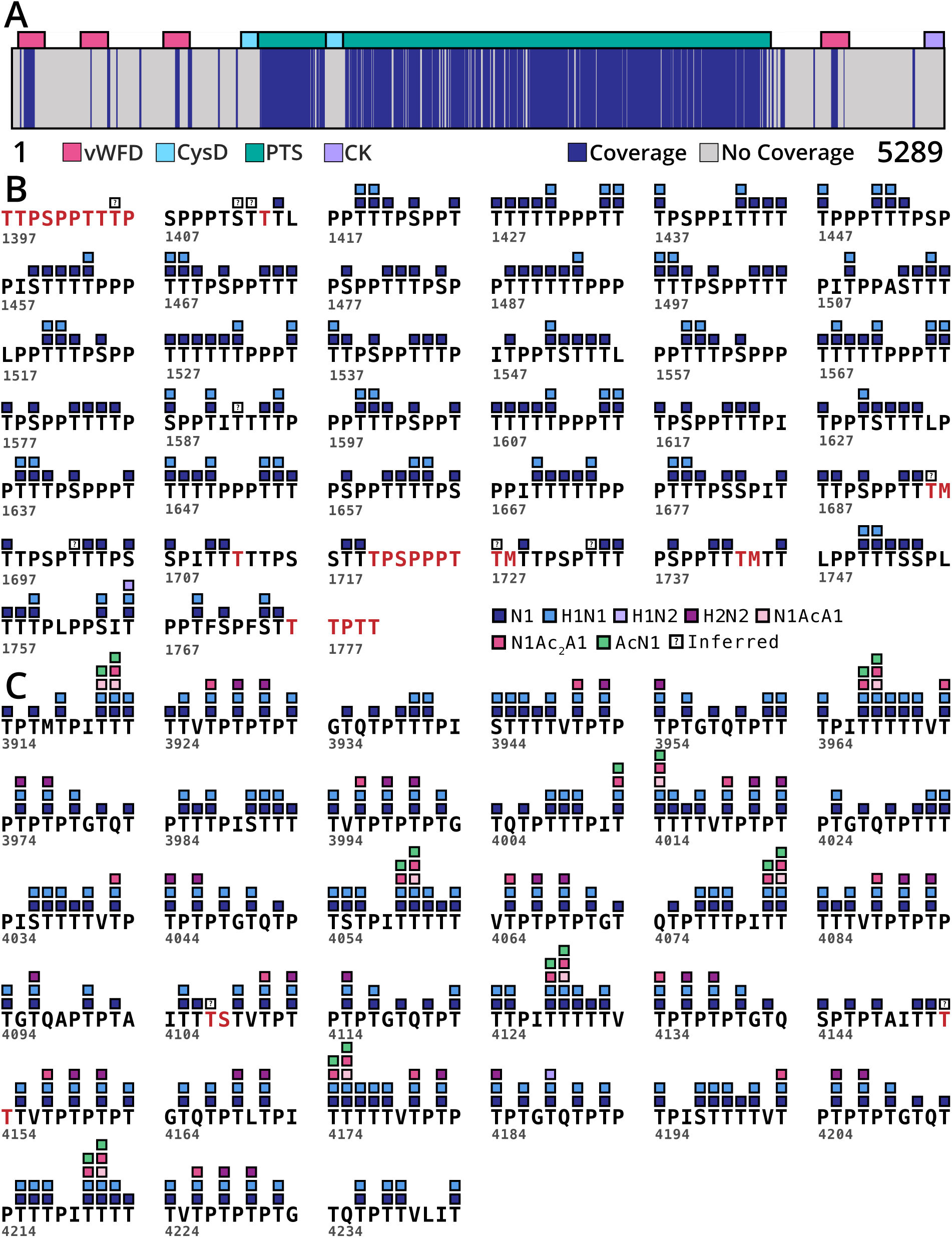
LC-MS analyses of Colon 1 & 2 yielded high sequence coverage of MUC2 and revealed domain-dependent glycosylation patterns. (A) Nearly complete coverage of MUC2’s mucin domains was obtained in LC-MS analyses, while proteomic coverage outside of the PTS regions was more limited. (B) Truncated O-glycans were confidently localized to residues in the first PTS domain, shown here in its entirety. (C) More varied O-glycan structures were localized to residues in the second PTS domain, a portion of which is shown here. “Inferred” indicates glycosites identified based on the cleavage preference of mucinase StcE. Red text indicates residues for which we did not obtain sequence coverage.

We suspect that protein structural differences between the two PTS domains could explain this disparity in glycosylation. Mucin domains are enriched in proline; in fact, mucins are proposed to have evolved from secreted proline-rich proteins in different animal lineages,^64^ underscoring the importance of this amino acid for mucin function. Most of the prolines in PTS 1 are found in clusters of two or three, while virtually all of the prolines in PTS 2 are separated by at least one other amino acid. The frequency and distribution of proline within the backbone has an enormous impact on protein tertiary structure and flexibility.^65^ For example, proline-rich repeat sequences are crucial for proper coiling of various collagens and their assembly into triple helices.^66^ The distinct proline distribution patterns in PTS 1 and PTS 2 therefore suggest that these domains adopt divergent structures. If so, the conformation of PTS 1 may sterically obstruct backbone association with Golgi glycosyltransferases that extend O-glycans beyond the core 1 structure. Our in-depth glycoproteomic mapping of a canonical mucin underscores the potential value of such analyses for revealing biological insights and potentially driving new research directions.

In all tumor samples, we identified glycopeptides decorated with a variety of O-glycan structures (**Figure S15a-d**). In the colorectal and esophageal tumor data, the distribution of identified glycoforms largely resembled that of MUC2 (**Figure S16a-c**). Notably, Neu5Ac-containing glycans were not particularly abundant in the colon and esophageal tumors, evidenced by traces of the ions for protonated Neu5Ac and its water loss at *m/z* 292.1027 and 274.0921, respectively (**Figure S17**). This result was surprising, as cancer is typically associated with hypersialylation of N- and O-glycans.^19^ We did detect some sialyl T-containing MUC2 glycopeptides in the colorectal tumors with LC-MS and MALDI-MSI, although at a much lower intensity than the un-sialylated versions of the same species (**Dataset S2, Figure S18**). If the tumors expressed high levels of Neu5Ac in the form of STn, a well-documented cancer antigen, it’s possible that StcE was unable to generate sialylated glycopeptides. Some data suggest that StcE does not readily cleave mucins heavily decorated with STn,^67^ however, the scarcity of ST and other Neu5Ac-containing structures in the MUC2 data suggests that StcE’s cleavage restrictions did not skew our glycoproteomic results. Due to limited sample availability, we were unable to confirm this hypothesis by staining tissues with sialic acid-binding immunoglobulin-like lectins (Siglecs) or plant-derived lectin probes.

Sialic acids are prone to in-source fragmentation (ISF) in both ESI and MALDI,^68^ which might have contributed to poor detection of sialylated glycopeptides. However, the Neu5Ac fingerprint signal was robust in the salivary gland tumor (**Figure S17**), thus we concluded that the unexpectedly low sialic acid content in other samples likely was not a technical artifact. Regardless, future studies may benefit from adding an on-slide derivatization step during sample preparation, which can reduce sialic acid lability during ionization and even distinguish its different glycosidic linkages.^69^ Possible biological explanations for the relative paucity of sialylated O-glycans in the gastrointestinal tumors should investigated in greater detail.

We explored possible modifications on Neu5Ac that could mask its presence in our data. Neu5Ac is often mono-, di-, or tri-acetylated in colon-derived glycans, and previous studies have linked loss of Neu5Ac acetylation with the development of colorectal cancer.^70^ In LC-MS analyses of both colon samples, we did detect low levels of mono- and di-O-acetylated Neu5Ac-containing glycans in both the tumor and non-tumor regions (**Figures S19-20**). Fingerprint ions for Ac_2_Neu5Ac were more intense than for AcNeu5Ac in both regions of Colon 2 as well as the non-tumor region of Colon 1 (**Figures S19-21**). However, the opposite pattern was detected in the tumor region of Colon 1 (**Figures S19-21**), although this observation may be driven by two peaks with particularly intense signal at retention times around 32 minutes (**Figure S19**). Glycans containing AcNeu5Ac and Ac_2_Neu5Ac could be localized on peptides from the second PTS domain of MUC2 (**Figures 4A & S22a-b, Dataset S2**). HCD spectra of species containing mono-acetylated Neu5Ac featured prominent marker ions at *m/z* 316.1026 and 334.1132. These fragments were detected in addition to *m/z* 358.1131 and 376.1237 in HCD spectra of glycopeptides containing Ac_2_Neu5Ac (**Figures 4A & S22a**). We were unable to detect glycopeptides decorated with AcNeu5Ac or Ac_2_Neu5Ac in MALDI-IMS experiments. Like sialic acids, acetyl groups are fairly labile, and are often lost prior to MS detection due to ISF.^71^ Failure to observe this species in MALDI-IMS analyses could be related to this technical challenge.

**Figure 4.**
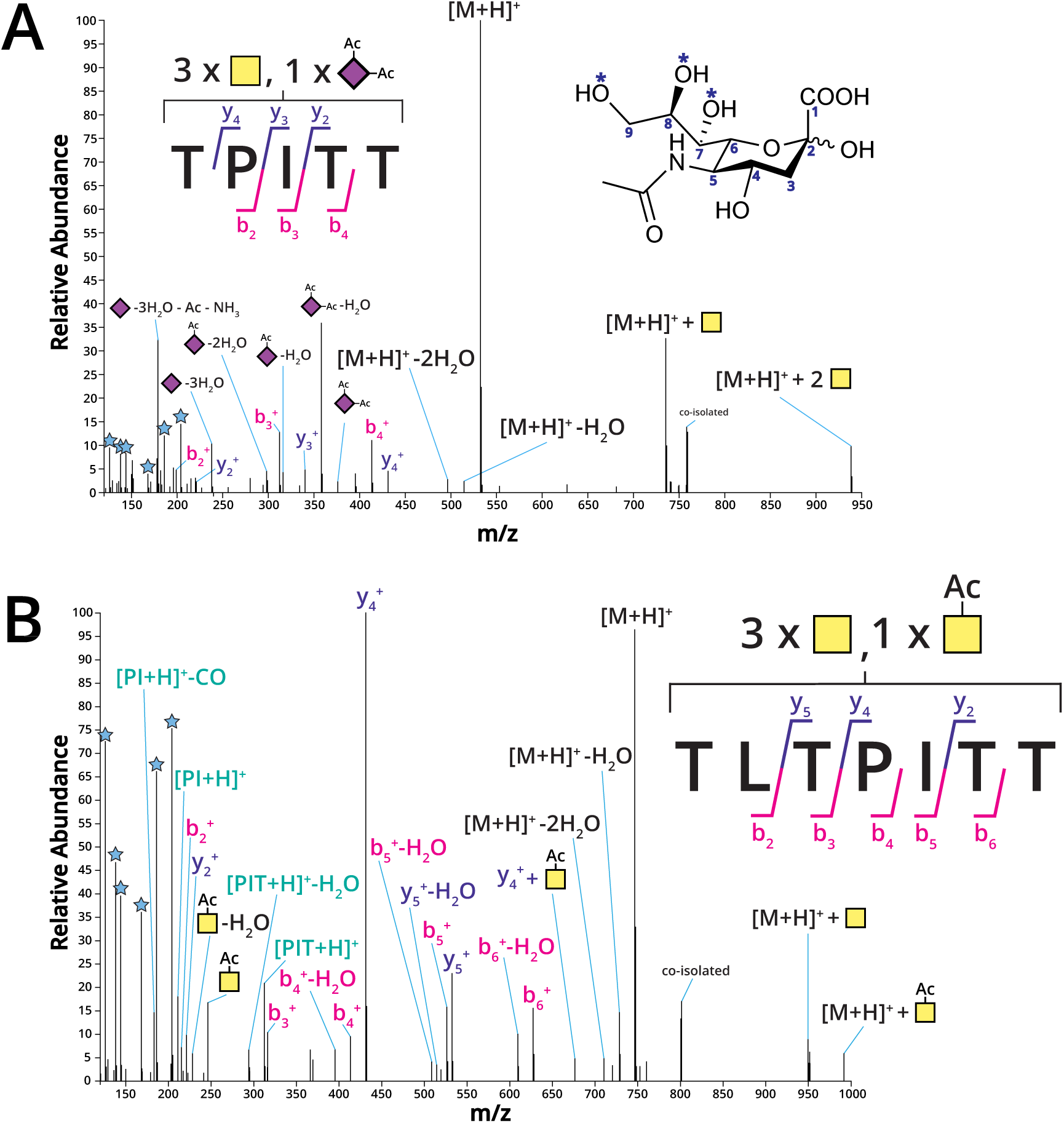
Spectral evidence for acetylated glycans on MUC2 sequences. (A) Representative higher-energy collision dissociation (HCD) spectrum for a glycoform of TPITT containing di-O-acetylated Neu5Ac. Marker ions for Ac_2_Neu5Ac are observed in the low-mass region, along with HexNAc fingerprint ions denoted by blue stars. [M+H]^+^ represents the mass of the naked peptide backbone. The inset shows the structure and numbered carbons of Neu5Ac, with asterisks indicating the most common sites of O-acetylation on the hydroxyl moieties of C7, C8, and/or C9. This specific spectrum is from Colon 2, region 2 data. (B) HCD spectrum for the sequence TLTPITT containing a putative AcGalNAc O-glycan. Fragments corresponding to the lone AcGalNAc as well as the y4+ and peptide backbone ions with AcGalNAc are observed. Internal fragments containing Pro, Ile, and Thr (from the sequence TPI or PIT) were also detected in the low-mass region.

### Glycoproteomic Analyses of Colorectal Tumors Reveal Acetylated GalNAc on MUC2

Some spectral evidence in both colon tumor samples suggested the presence of an additional acetyl moiety on GalNAc (**Figures 4B & S23a-b**). This included fragments in HCD spectra corresponding to the HexNAc fingerprint ions at *m/z* 204.0867 and *m/z* 186.0761 shifted by the mass of an acetyl group (+42.0106 Da), resulting in glycan fragments with theoretical *m/z* 246.0973 and 228.0867 (**Figures 4B & S23a**). For all glycopeptides identified with this modification, the naked peptide backbone was detected at the expected *m/z* in HCD spectra. This observation ruled out potential acetylation of the peptidyl content of the glycopeptide, adding additional support for acetylation on one of the glycans. We detected LC-MS evidence for “AcGalNAc” only in the colorectal tumor regions. MALDI-IMS results for Colon 1 included a heat map for an *m/z* corresponding to the mass of TVTPTPTPTG decorated with four Tn antigens and one AcGalNAc, the presence of which was identified by LC-MS (**Figure S24a, Dataset S2**). Although the MALDI-IMS intensity of this ion was very weak, its distribution overlapped with the expression of Tn-only MUC2 glycopeptides in the tumor region (**Figure S24a-b**). While the amine nitrogen in GalNAc could theoretically accommodate a second acetyl group, we suspect the additional acetylation is O-linked through one of the hydroxyl moieties, similar to sialic acid acetylation. Although others have speculated about its existence, to our knowledge, experimental evidence for endogenous O-acetylation of GalNAc in O-glycans has not been reported in the literature. Additional studies should more thoroughly assess whether AcGalNAc is a true biological phenomenon or a technical artifact.

### LC-MS Detects MUC2 Glycoforms and Glycosylated Dipeptides in Esophageal Tumor

Numerous tumor-associated ions were detected in the esophageal tissue by MALDI-MSI, but we were unable to identify any of these species in the associated LC-MS data (**Figure S25**). Conversely, although many MUC2 O-glycopeptides were identified in the esophageal tumor LC-MS data, we were unable to detect any by MALDI-MSI. This tumor was notably smaller than those in the colorectal and salivary gland tissues, and as a result, the amount of recovered material and subsequent LC-MS HexNAc signal was lower in the esophageal samples compared to the other tumors (**Figure S8a**). Many glycopeptide species were too weak for MS2 selection and were identified based on precursor mass and retention time (**Dataset S2**). The most abundant MUC2 glycopeptides were decorated only with the Tn antigen, a particularly interesting result if this tumor is indeed a metastasis, as described above. Previous work using an *in vitro* skin model showed that cancer cells with O-glycosylation limited to the Tn and STn antigens exhibited a significantly more invasive phenotype compared to cells decorated with wildtype O-glycosylation.^20^

Despite strong HexNAc ion signal in the esophageal tumor (**Figure S8a**), relatively few spectra could be confidently assigned to glycopeptides from specific proteins. Dipeptides with the sequence TI/TL or IT/LT were highly abundant and decorated with a variety of structures, including H2N2F1 and H2N3F2 (**Figure S26a-b)**. The latter of these may correspond to either the Lewis b or Lewis y blood group antigens, which are indistinguishable with our MS/MS acquisition method. While these species were most likely cleaved from MUC2, we cannot confidently confirm this due to the lack of unique peptide sequence. Additionally, much of the HexNAc ion signal could be attributed to free N-glycans released during PNGaseF treatment (**Figure S27a**). HCD spectra of free N-glycans included the chitobiose marker ion at m/z 893.3250, and the relative intensity of the HexNAc marker ion at m/z 138.055 was much higher than the m/z 144.0655 ion, indicative of high GlcNAc content (**Figure S27b**).

While free N-glycans and O-glycosylated dipeptides could be detected in other samples, their relative abundance compared to the overall HexNAc signal was particularly high in the esophageal tumor. This result could stem from the small size of the esophageal tumor or lower endogenous mucin production, resulting in only a fraction of the glycopeptide content of the other samples. Another possibility is that the mucins produced by this tumor were largely resistant to StcE digestion. Due to limited sample availability, we did not perform additional staining or glycoproteomic experiments to further investigate these results.

### Versican Dominates the Glycoproteome of Salivary Gland Tumor

In the salivary gland tumor, glycopeptide results were dominated by the core protein of versican, a mucinous chondroitin sulfate proteoglycan (**Figure S16d, Dataset S2**). A few versican glycopeptides were also detected in other samples, albeit very few at relatively low abundances (**Dataset S2**). Various cancers are associated with versican overexpression,^72^ and cleavage of the core protein by tissue-resident and tumor-associated proteases may contribute to tissue invasion and metastasis.^73^ Core 1 and core 2 structures with and without sialylation were localized on versican glycopeptides (**Figure S15d**), although the majority were decorated with only T antigens (**Figure S14b**). Surprisingly, very few versican glycopeptides displayed the Tn antigen, in stark contrast to MUC2 results in the other tumors (**Figure S14a-b**). Interestingly, many versican peptide backbone sequences corresponded to non-StcE proteolytic cleavage at the N- or C-terminus (**Dataset S2**). We detected only low levels of non-specific cleavage in other samples, suggesting degradation of versican by endogenous enzymes prior to tissue resection and preservation. Unmodified peptide searches identified multiple endogenous proteases and peptidases in this tissue (**Table S2**). This includes matrix metalloprotease 9 (MMP-9), also known as gelatinase B, an ECM modulator that is strongly implicated in metastatic invasion.^74^ A previous study demonstrated MMP-9 readily degraded versican extracted from rabbit lung, although the exact cleavage sites were not determined.^75^ MMP-9 was also identified in unmodified peptide data for the non-tumor region of Colon 1, both regions in Colon 2, and the esophageal tumor (**Table S2**). Whether O-glycosylation influences versican proteolysis and tumor ECM remodeling should be explored in greater detail.

## Concluding Remarks

Our results demonstrate the potential of our glycopeptide imaging workflow for studying the tumor microenvironment, but important limitations of this approach should be considered in future efforts. One of the major technical limitations of this approach was the relatively low sensitivity of MALDI-IMS compared to LC-MS, as discussed above. Furthermore, it is possible that the tumor molecular profile varied by tissue slice, which might have contributed to differences in the species detected by LC-MS and MALDI-MSI. Another drawback of our approach is that isobaric MUC2 glycopeptides detected by LC-MS could not be distinguished by MALDI-MSI, as MS2 spectra were not acquired with this instrument. For example, LC-MS analyses identified glycoforms of TTPSPPTI, TPSPPITT, TPITPPTS, and TVTPTPTP, all of which have the same theoretical monoisotopic mass. Lastly, although we searched for numerous O-glycan structures in our LC-MS analyses, it is possible that we failed to identify glycopeptides decorated with relatively uncommon structures or modifications. Future work would ideally include a separate glycomics analysis of each tissue. This data could be used to generate a tailored glycan database to be used in glycoproteomic searches, similar to how we constructed custom protein databases by analyzing unmodified tryptic peptides.

In summary, we developed a coupled-MS workflow to visualize specific mucin glycoforms in tissue. Our approach also enabled in-depth glycoproteomic mapping of MUC2, with near-complete sequence coverage over its two PTS domains for the first time. Although most of the localized O-glycans were the Tn or T antigen, we also identified core 2 structures, mono- and di-acetylated STn-containing glycans, and potentially acetylated GalNAc. In the future, additional glycoepitopes may be revealed by incorporating alternative mucinases into our workflow. For example, the cleavage motif of SmE, recently characterized by our lab,^10^ is significantly less restricted than that of StcE. We also hope to assess the relative frequency of the distinct Tn to T antigen expression patterns detected in the colon samples by significantly expanding our patient cohort. Despite the strong link between these structures and numerous cancers, malignant processes driven by the Tn and T antigens are still poorly understood at the mechanistic level. Further interrogation of Tn and T expression patterns in a variety of clinical specimens may provide useful insights and drive new biological hypotheses in cancer research. Overall, we present a powerful method to image and sequence cancer-associated mucin glycopeptides *in situ*. We anticipate that this workflow can be applied to many cancers, tissues, and/or pathologies and will prove useful to the broader glycobiology, cancer biology, and mass spectrometry fields.

## Materials and Methods

### Tissue Acquisition, Fixation, and Sectioning

A total of four archival formalin-fixed, paraffin-embedded (FFPE) carcinoma tissue blocks (Colon 1, Colon 2, Esophagus, and Salivary Gland) were obtained from the Medical University of South Carolina’s Hollings Cancer Center Biorepository. No identifiable patient information was provided to the investigators, and the Institutional Review Board at the Medical University of South Carolina approved the use of all tissues (protocol 17669). Colon 1 and Colon 2 were derived from the same donor, while the Esophagus and Salivary Gland samples were obtained from two additional patients. A pathologist annotated the boundaries of the tumor and adjacent non-malignant regions for each. Tissues were sectioned onto slides at thickness of 5 µM for further sample preparation.

### Tissue Preparation for MS Experiments

His-tagged mucinase StcE was expressed and purified as previously described^48^ using a pET28 plasmid kindly provided by the Bertozzi laboratory. All solvents for slide preparation were purchased from Fisher Scientific. Tissue slides were prepared as described previously.^76^ Slides were heated, tissue side-up, at 60 °C for 1 hour. After dewaxing and rehydration of tissue slides, antigen retrieval was performed in citraconic anhydride buffer, pH 3, for 30 minutes in a decloaking chamber set to 95 °C. Following a water wash, slides were dried in a desiccator. A total of 15 passes of 0.1 µg/µL PNGaseF PRIME (N-Zymes Scientific, Doylestown, PA) was applied to slides for N-glycan release using an M5 Sprayer (HTX Technologies, Chapel Hill, NC) with the following parameters: 25 µL/min flow rate, 1200 mm/min velocity, 3-mm offset, 10 psi, and 45 °C. Deglycosylation in prewarmed humidity chambers proceeded for 2 hours at 37 °C, after which tissues were H&E stained and scanned. Cover slips were removed, then washed sequentially in 10 mM Tris pH 8.5 and citraconic anhydride pH 3 buffers, then water to remove stain and released N-glycans, and dried completely in a desiccator. The M5 sprayer then applied mucinase StcE using the same parameters as above. StcE digestion in the same prewarmed humidity chambers proceeded overnight at 37 °C and slides were dried in a desiccator prior to further preparation for MALDI-MSI or LC-MS analyses.

### MALDI-MSI Matrix Application

A matrix of 7 mg/mL CHCA in 50%ACN/0.1% TFA was applied to slides with the M5 sprayer at 10 psi and 79 °C using a flow rate of 100 µL/min, velocity of 1300 mm/min, and a 2.5-mm offset. Slides were stored in a desiccator until MALDI-MSI acquisition.

### MALDI-MSI Data Acquisition and Processing

All samples except Colon 2 were analyzed with a timsTOF fleX MALDI-QTOF mass spectrometer (Bruker Corporation, Billerica, MA). The following parameters were used: mass range 300-3500 m/z, positive ion mode, 20-25 µm laser spots, 300 shots per pixel, and 40-µm raster. Colon 2 was analyzed with a SolariX dual-source 7T MALDI-FTICR mass spectrometer (Bruker). Settings were as follows: mass range 300-3500 m/z, positive ion mode, 20-25 µm laser spots, 300 shots per pixel, and 40-µm raster. Raw imaging data was processed in SCiLS Lab version 2024b Pro (Bruker Corporation, Billerica, MA), and O-glycopeptide spectra were normalized to total ion count.

### Additional Mucinase Digestion and Glycopeptide Extraction for LC-MS

LC-MS grade water (Thermo Scientific, 51140), acetonitrile (ACN) (Honeywell, LC015), methanol (MeOH) (Fisher Scientific, A456), and formic acid (FA) (Thermo Scientific, 85178) were used unless otherwise noted. Slide humidity chambers were prepared by placing dish sponges saturated with Milli-Q water (MicroPure UV/UF, Thermo Scientific, 50132370) in the bottom of airtight plastic containers, then pre-warming at 37 °C for at least 1 hour. The tumor and adjacent non-tumor tissue regions were outlined using a hydrophobic barrier pen (Sigma Aldrich, Z377821 or Z672548), then dried for approximately 5 minutes. A 0.05 mg/mL solution of StcE in fresh, 50 mM ammonium bicarbonate (AmBic) (Honeywell Fluka, 40867) was deposited via micropipette to completely cover each outlined region. Slides were placed in pre-warmed humidity chambers and incubated at 37 °C for 13-23 hours.

After removing slides from the humidity chambers, remaining solution from each region was collected via pipette into separate 1.5 mL Eppendorf tubes, taking care not to touch tissue surfaces with pipette tips. Glycopeptides were collected into the same tubes after extraction with 3 different solvents: 0.1% FA in water, 4:1 ACN:0.1% FA in water, and 7:3 MeOH:0.1% FA in water.^42^ Using the same volume as StcE digestion, each solvent (in the order listed) was pipetted up and down ten times on the surface of the slide. This process was performed twice for each of the solvents, for a total of six extractions.

After glycopeptide extraction, slides were allowed to dry in a chemical fume hood. A 0.01 mg/mL solution of sequencing-grade trypsin (Promega, V5113) in 50 mM AmBic was applied to slides via micropipette, using the same volumes as the mucinase digestion. Slides were placed in humidity chambers and incubated at 37 °C for 4-5 hours before removing tryptic peptides using the extraction method described above. All samples were taken to dryness in a vacuum centrifuge and stored at -20 °C until further use.

### Glycopeptide Sample Preparation for LC-MS

Glycopeptide samples were resuspended in 100 µL of 50 mM ammonium bicarbonate, after which the approximate protein concentration was measured by NanoDrop One Microvolume UV-Vis Spectrophotometer (Thermo-Fisher). Samples were acidified with 1 μL of neat FA. Desalting was performed using 10 mg Strata-X 33 µm polymeric reversed phase SPE columns (Phenomenex, 8B-S100-AAK). Each column was activated using 1 mL of ACN, equilibrated with 1 mL of 0.1% FA, pre-eluted with 1 mL of 0.1% FA in 40% ACN, and then re-equilibrated with 1 mL of 0.1% FA. Samples were added to the column and de-salted using two 150-µL rinses of 0.1% FA. The columns were transferred to clean 1.5 mL tubes and peptides were eluted with two 150 µL additions of 0.1% FA in 40% ACN. Samples were taken to dryness in a vacuum centrifuge (LabConco), resuspended in 7-10 µL of 0.1% FA, and transferred to HPLC vials for LC-MS analysis.

### Tryptic Peptide Sample Fractionation and Preparation for LC-MS

Tryptic peptide samples were resuspended in 100 μL of 50 mM ammonium bicarbonate, after which the approximate protein concentration was obtained via NanoDrop One. Samples were acidified by adding 1 μL of neat FA, desalted as above, and taken to dryness.

To obtain deeper proteome coverage, peptides were fractionated with hydrophilic interaction chromatography (HILIC)-based solid-phase extraction (SPE) using 12-μm, 300-Å PolyHYDROXYETHYL Aspartamide™ (PHEA) (PolyLC®, BMHY12-03). We constructed PHEA-packed SPE columns based on the stop-and-go extraction tips (StageTips) design.^77^ Briefly, a 16-gauge blunt-tip needle (Cadence Science, 7938) was used to punch glass filter membranes from a larger disk (Whatman, grade GF/F, 987468). Using a 20-mL plastic syringe (Henke Sass Wolf, Henke-Ject® Luer Lock, 4200.X00V0), the glass membranes were then plugged into the ends of 200-μL micropipette tips, one for each sample. The plugged tips were then placed in 1.5-mL tubes equipped with centrifuge adapters (GL Sciences, 5010-21514).

A small scoop of PHEA resin was resuspended in 0.5 mL of 200 mM ammonium formate (AF) (Alfa Aesar, 14517) in a 1.5-mL tube, and equal amounts of slurry were transferred to each packed pipette tip. PHEA resin was packed into beds of approximately 1 cm in length by centrifugation at 350-400 rcf. All subsequent centrifugation steps used the same speed. Packed tips were rinsed with 100-uL volumes of the following solvents: 1% FA, 20 mM AF in 90% ACN, 200 mM AF, pure water, 20 mM AF in 90% ACN, and a final wash with 20 mM AF in 90% ACN. Dried peptide samples were resuspended in 200 mM AF to a final concentration of 1 mg/mL. A 10-μL aliquot was transferred to a tube containing 90 μL of neat ACN, producing 10-μg peptide aliquots in 20 mM AF in 90% ACN.

Samples were applied to packed tips, then rinsed with two 100-μL volumes of 20 mM AF in 90% ACN. The packed tips were transferred to clean tubes, and less hydrophilic peptides were eluted with two 100-μL volumes of 40 mM AF in 80% ACN (‘Fraction 1’). The packed tips were transferred again to clean tubes, and more hydrophilic peptides were eluted with 150 μL, then 100 μL of 1% FA (‘Fraction 2’). This final elution was performed at 200 rcf. Both eluted fractions were taken to dryness, after which samples were resuspended in 7-10 µL of 0.1% FA in water and transferred to HPLC vials for LC-MS analysis.

### LC-MS Data Acquisition

Samples were analyzed by online nanoflow liquid chromatography-tandem mass spectrometry using an Orbitrap Eclipse Tribrid mass spectrometer (Thermo Fisher Scientific) coupled to a Dionex UltiMate 3000 HPLC (Thermo Fisher Scientific). A volume of 6 µL for tumor glycopeptide samples or 1-6 µL for PHEA-enriched fractions was injected onto an Acclaim PepMap 100 column packed with 2 cm of 5 µm C18 material (Thermo Fisher, 164564) using 0.1% FA in water (solvent A). Peptides were then separated on a 15 cm PepMap RSLC EASY-Spray C18 column packed with 2 µm of C18 material (Thermo Fisher, ES904) using a gradient from 0-35% solvent B (0.1% FA with 80% ACN) in 60 min.

For glycopeptide and PHEA-enriched Fraction 2 samples (as well as Fraction 1 for Colon 2 and the Salivary Gland), full scan MS1 spectra were collected at a resolution of 60,000, an automatic gain control (AGC) target of 3e5, and a mass range from 400-1500 m/z or 300-1500 m/z. Charge states 2-6 or 2-7 were selected for fragmentation. MS2s were generated at top speed for 3 seconds. Higher-energy collisional dissociation (HCD) was performed on all selected precursor masses with the following parameters: isolation window of 2 m/z, stepped collision fragmentation at 25-30-35% or 25-30-40% normalized collision energy (nCE), Orbitrap detection (resolution of 7,500), maximum inject time of 75 ms, and a standard AGC target. Dynamic exclusion was enabled with a repeat count of 2, repeat duration of 8 s, and exclusion duration of 8 s.

Detection of at least 3 HexNAc or Neu5Ac fingerprint ions (126.055, 138.055, 144.07, 168.065, 186.076, 204.086, 274.092, 292.103, and 316.102) present at ± 10 ppm and greater than 5% relative intensity triggered additional fragmentation on precursors below 850 m/z (with an intensity of at least 5e4) using electron transfer dissociation (ETD), and fragments were detected in the ion trap at a normal scan rate. For precursors between 850-1300 m/z (with an intensity over 5e5), detection of fingerprint ions triggered ETD with supplemental activation (EThcD) at 15% nCE, which was analyzed in the Orbitrap with a resolution of 7,500. Calibrated charge-dependent reaction times were used for ETD and EThcD fragmentation, with 200 ms maximum injection time and custom injection targets. Dynamic exclusion was enabled with a repeat count of 2, repeat duration of 10 s, and exclusion duration of 10 s.

For PHEA-enriched Fraction 1 samples of Colon 1 and the Esophagus, full scan MS1 spectra were collected at a resolution of 60,000, an automatic gain control (AGC) target of 3e5, and a mass range from 400-1500 m/z or 300-1500 m/z. Charge states 2-6 were selected for fragmentation. MS2s were generated at top speed for 3 seconds. Higher-energy collisional dissociation (HCD) was performed on all selected precursor masses with the following parameters: isolation window of 2 m/z, stepped collision fragmentation at 27-29-31% or 25-30-35% normalized collision energy (nCE), Orbitrap detection (resolution of 15,000), maximum inject time of 100 ms, and a standard AGC target. Dynamic exclusion was enabled with a repeat count of 2, repeat duration of 7 s, and exclusion duration of 8 s. No electron-based MS2 spectra were collected for these samples.

### Unmodified Proteome Searches of PHEA-enriched Tryptic Peptides

Tryptic peptide *.raw files were searched against the human proteome with Byonic (Protein Metrics). Mass tolerances of 10 ppm, 20 ppm, and 0.3 Da were used for MS1, FTMS MS2, and ITMS MS2, respectively. Met oxidation was selected as a variable modification. The search was set to “fully specific” with proteolytic cleavage C-terminal to R and K, allowing 2 missed cleavages.

### Glycoproteomic searches in O-Pair

Glycopeptide *.raw files for each tumor region were searched using O-Pair with MetaMorpheus (v1.0.2 or v1.0.3), using a manually curated protein database containing mucins of interest and StcE. The dissociation type was set to HCD, and EThcD was selected for child scan dissociation. Precursor and product ion mass tolerances were set to 10 ppm and 20 ppm, respectively. “Non-specific” protease cleavage was selected, and peptide length was set to 5-25 amino acids. A maximum of 6 O-glycans were allowed, using the “OGlycan.gdb” database. All glycopeptide identifications assigned Level 1 or 1b were validated by manual inspection of raw spectra using QualBrowser in XCalibur (v4.1.31.9).

### Assignment of Identified Glycopeptides to MALDI-MSI Spectra

Subsequently, putative LC-MS *m/z* values corresponding to doubly and triply protonated precursor ions were calculated for each *m/z* detected by MALDI-MSI. Raw LC-MS data were then searched manually for the possible precursor masses. If one of the calculated *m/z* values was detected, the O-Pair output file was checked for any identifications with the *m/z* detected. If there were no O-Pair identifications for a detected *m/z*, the corresponding HCD and MS2 spectra were used for manual peptide sequencing and glycan localization. Sequenced peptides were then matched to specific proteins using NCBI’s BLAST algorithm.

### Data Availability

The LC-MS proteomics raw data and search outputs have been deposited to the ProteomeXchange Consortium via the PRIDE^78^ partner repository with the dataset identifier PXD055865.

## Acknowledgments

S.C.L. is supported by the National Institutes of Health Chemical Biology Training Grant (T32 GM067543); K.E.M. is supported by the Yale Endowed Postdoctoral Fellowship in the Biological Sciences. S.A.M. is supported by the Yale Science Development Fund and NIGMS R35-GM147039. R.R.D. is supported by R33CA267226, U01CA242096, P30DK123704 and the Biorepository and Tissue Analysis Shared Resource, Hollings Cancer Center, Medical University of South Carolina (P30 CA138313). V.V. is supported by NIH/NIAAA 5R24AA022057. We thank Carolyn Bertozzi (Stanford University) for the StcE plasmid, and fellow Malaker lab members Joann Chongsaritsinsuk and Vincent Chang for expressing and purifying the StcE used in this project. We thank Dr. Gunnar Hansson for helpful conversations regarding acetylation of Neu5Ac and GalNAc.

