## Supplemental Information for "Glycosite Mapping and *in situ* Mass Spectrometry Imaging of MUC2 Glycopeptides via On-slide Digestion with Mucinase StcE"

### Colon 1

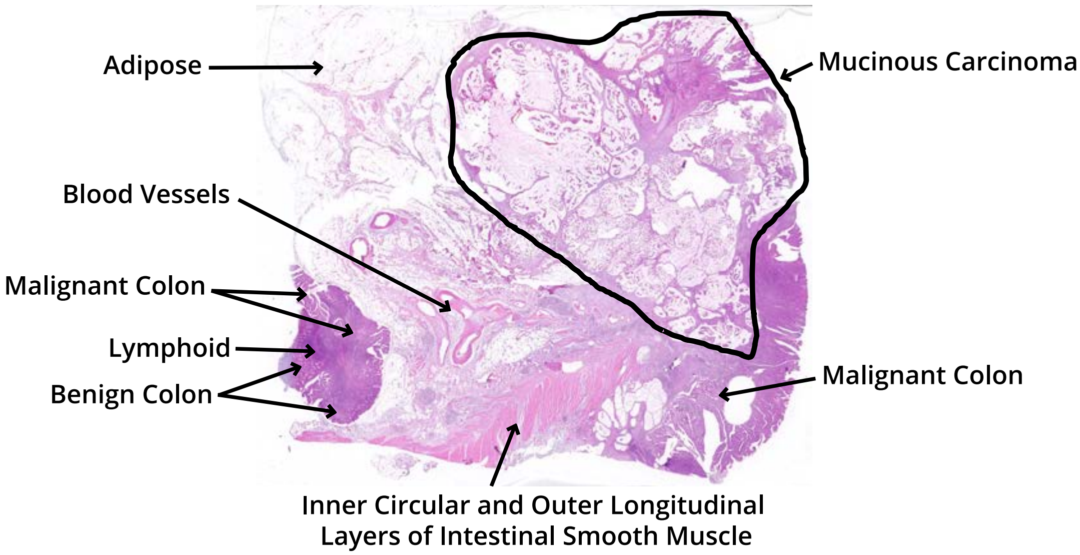


**Fig. S1.** H&E stain of Colon 1, with tissue features annotated and tumor region encircled. Tumor infiltration into muscle tissue is observed.

### Colon 2

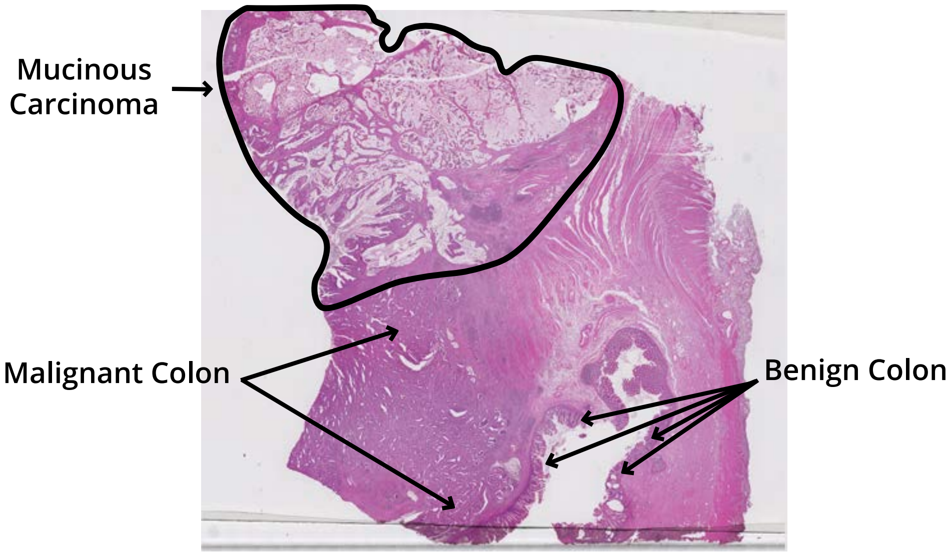


**Fig. S2.** H&E stain of Colon 2, with tissue features annotated and tumor region encircled. This tumor was resected from the same patient as Colon 1. Tumor infiltration into muscle tissue is observed.

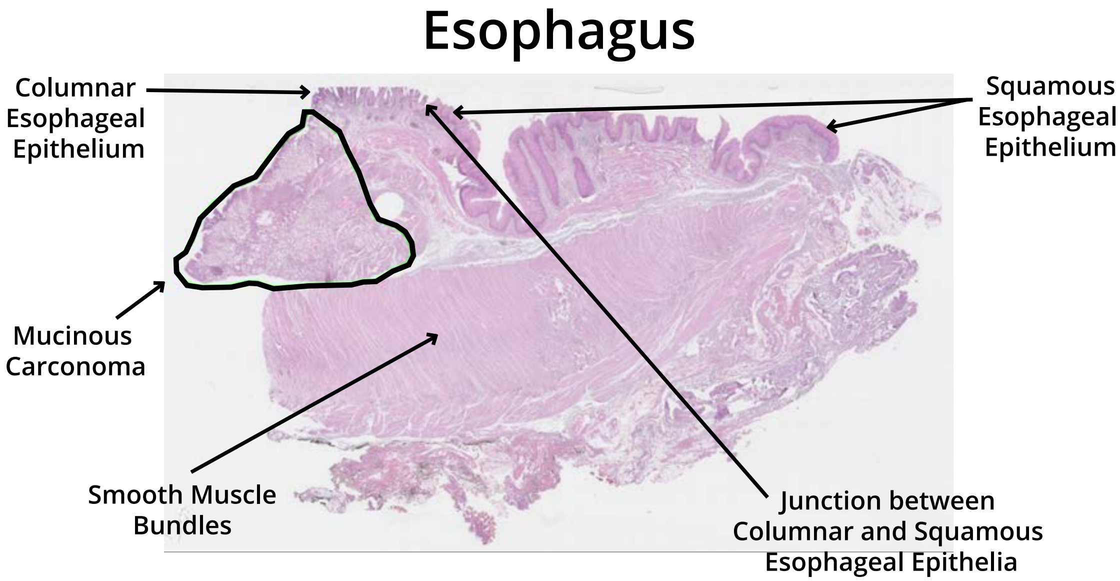


**Fig. S3.** H&E stain of the esophageal sample, with tissue features annotated and tumor region encircled.

### Salivary Gland

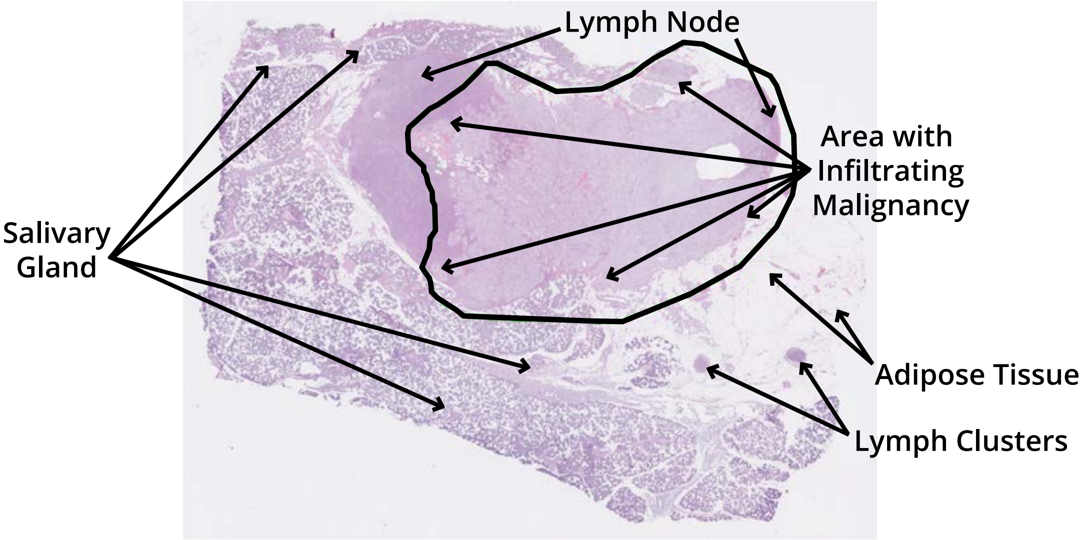


**Fig. S4.** H&E stain of the salivary gland sample, with tissue features annotated and tumor region encircled.

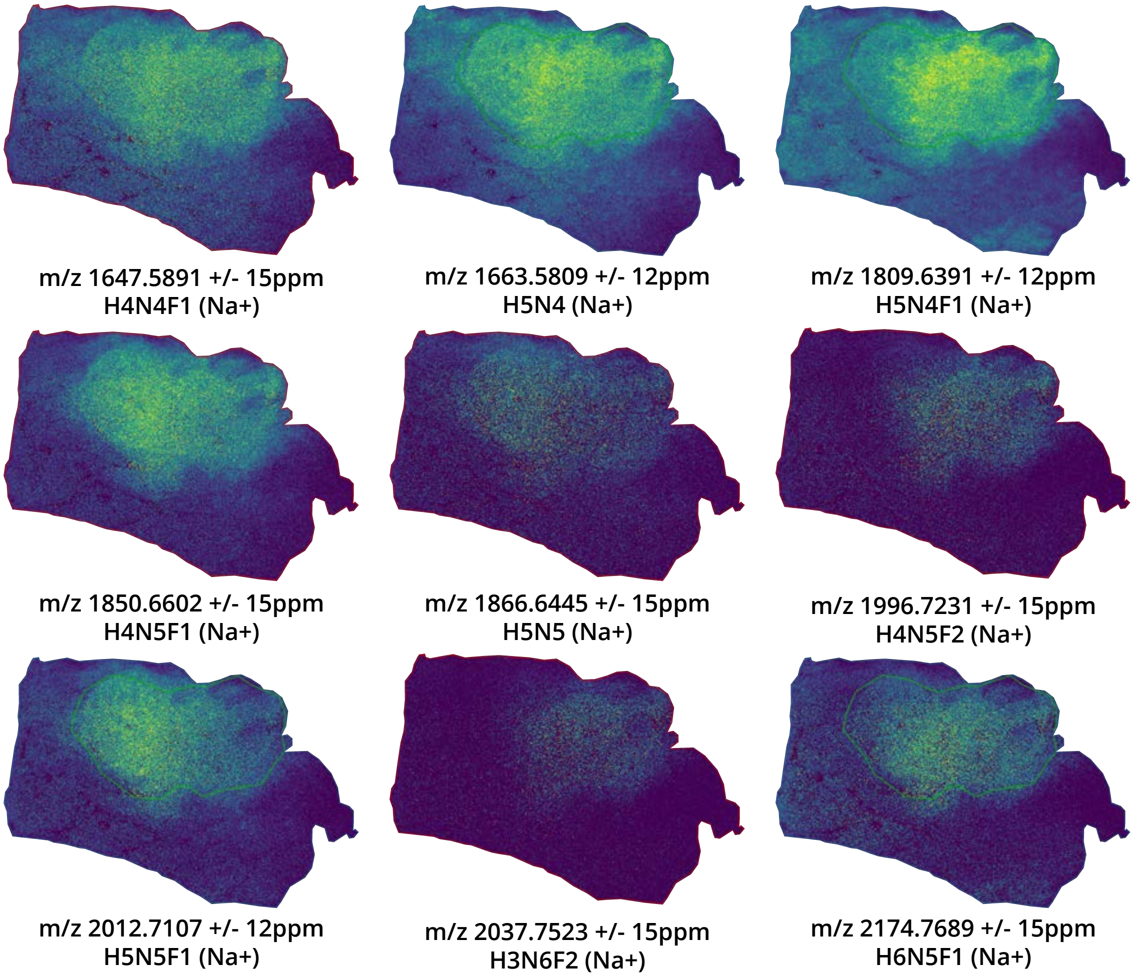


**Fig. S5.** MALDI-MSI images of N-glycans detected within the boundaries of the salivary gland tumor treated with PNGaseF and mucinase StcE. $\alpha$-cyano-4-hydroxycinnaminic acid (CHCA) matrix was applied via automatic sprayer prior to acquisition with a timsTOF fleX MALDI-QTOF mass spectrometer (Bruker). Images were generated with SCiLS Lab 2024b Pro (Bruker) by manually selecting m/z values within the mass window indicated. All identified species correspond to complex-type N-glycans, and all depicted masses correspond to mono-sodiated precursor ions. were mono-sodiated.

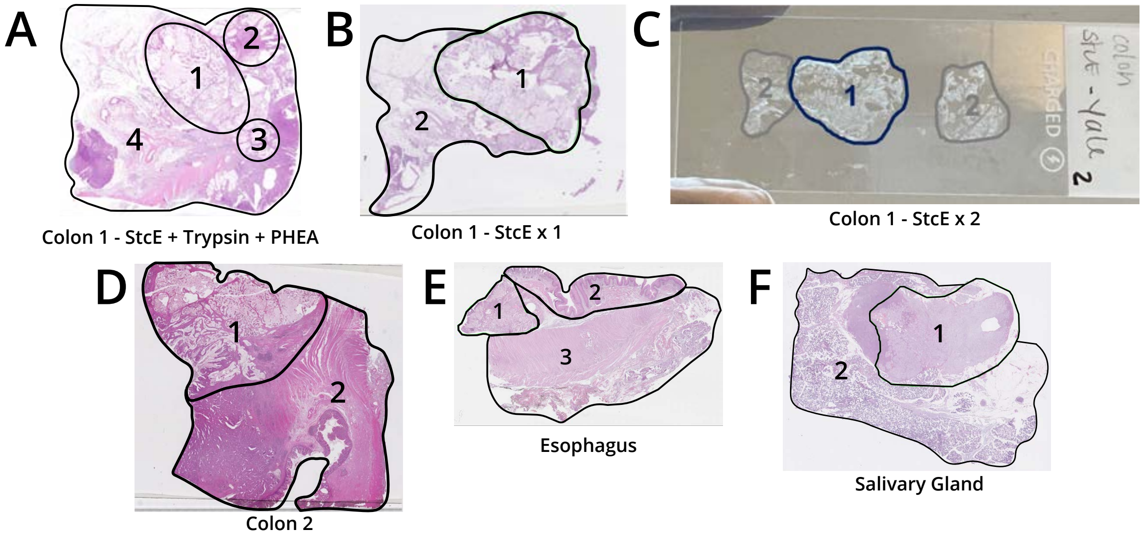


**Fig. S6.** Tumor and non-tumor regions were outlined with a hydrophobic barrier pen for LC-MS experiments. (A) Based on intratumor expression patterns observed in MALDI-MSI, initial experiments with Colon 1 divided the tumor into Regions 1, 2, and 3, while the rest of the tissue was designated Region 4. (B) A different tissue section of Colon 1 used for a StcE-only experiment. Regions 1 and 2 correspond to the tumor and non-tumor regions, respectively. (C) A third section of Colon 1 used for double-StcE experiments. Regions 1 and 2 correspond to the tumor and non-tumor regions, respectively. The separate non-tumor regions were combined into a single sample for this particular tissue. (D) Colon 2 was separated into Regions 1 (tumor) and 2 (non-tumor). (E) In the esophageal sample, the tumor was designated Region 1. Because we anticipated high mucin content in the villous tumor-adjacent area, we separated Region 2 from the rest of the non-tumor Region 3. (F) The salivary gland sample was separated into Regions 1 (tumor) and 2 (non-tumor).

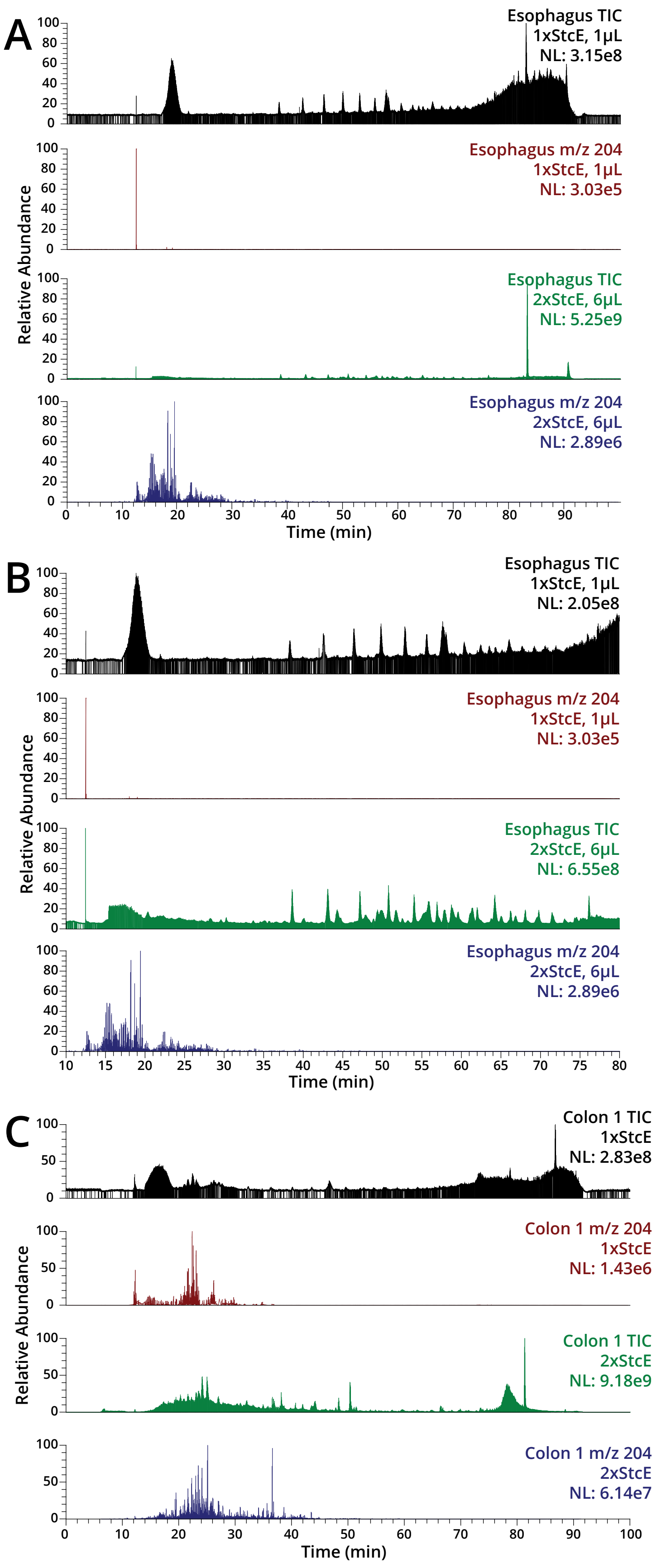
A
Esophagus TIC
1xStcE, 1μL
NL: 3.15e8
Esophagus m/z 204
1xStcE, 1μL
NL: 3.03e5
Esophagus TIC
2xStcE, 6μL
NL: 5.25e9
Esophagus m/z 204
2xStcE, 6μL
NL: 2.89e6
Relative Abundance
Time (min)
B
Esophagus TIC
1xStcE, 1μL
NL: 2.05e8
Esophagus m/z 204
1xStcE, 1μL
NL: 3.03e5
Esophagus TIC
2xStcE, 6μL
NL: 6.55e8
Esophagus m/z 204
2xStcE, 6μL
NL: 2.89e6
Relative Abundance
Time (min)
C
Colon 1 TIC
1xStcE
NL: 2.83e8
Colon 1 m/z 204
1xStcE
NL: 1.43e6
Colon 1 TIC
2xStcE
NL: 9.18e9
Colon 1 m/z 204
2xStcE
NL: 6.14e7
Relative Abundance
Time (min)

**Fig. S7.** Glycopeptide signal comparison for one or two StcE treatments. (A) Total ion chromatograms (TICs) and traces of HexNAc fingerprint ion at *m/z* 204.0867 for the esophageal tumor digested with StcE once (black, red) or twice (green, blue). (B) Zoom-in on the same TICs and HexNAc traces from 10-80 minutes. (C) Total ion chromatograms (TICs) and traces of HexNAc fingerprint ion at *m/z* 204.0867 for Colon 1 tumor digested with StcE once (black, red) or twice (green, blue).

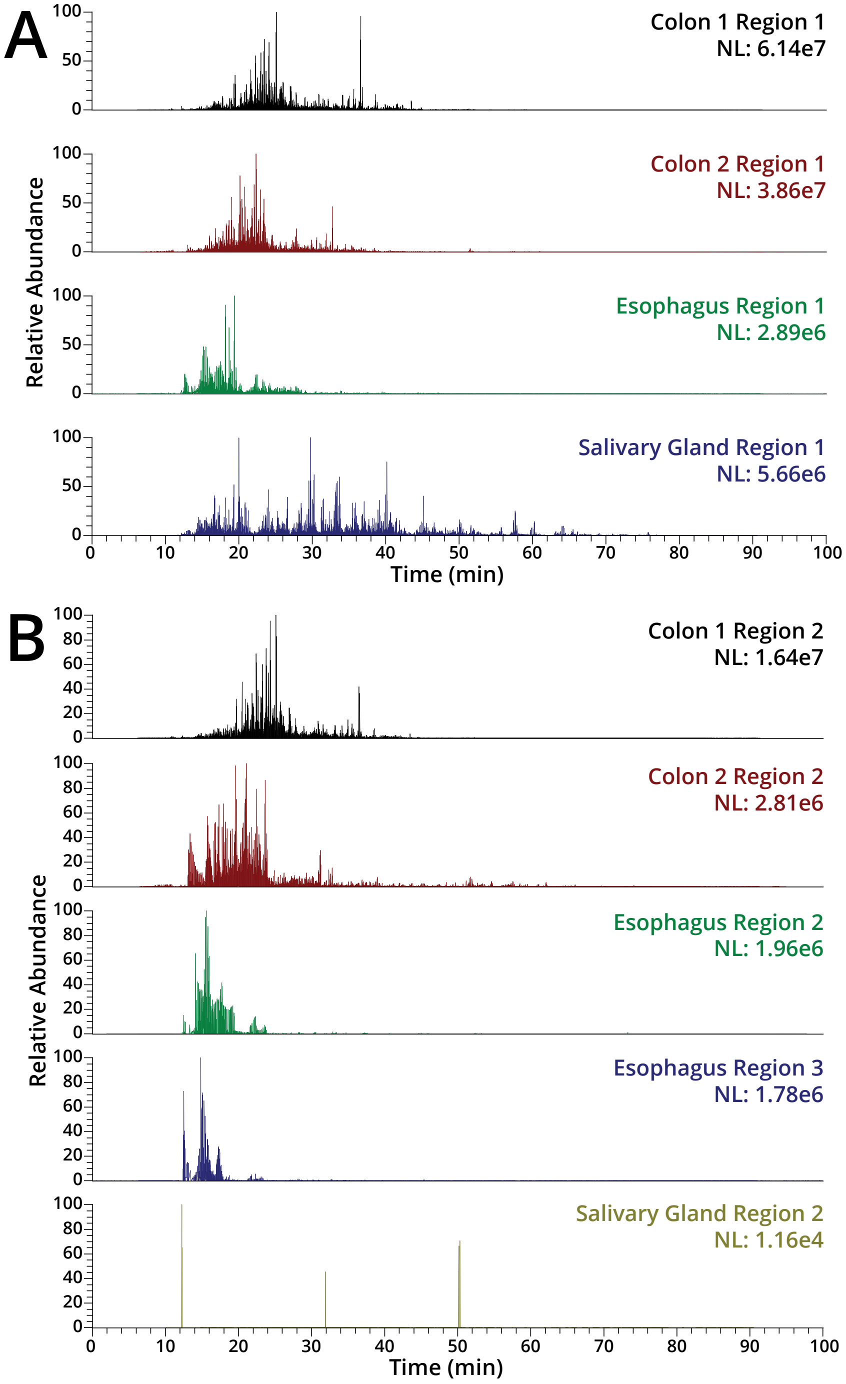


**Fig. S8.** Saturating the tissue surface with StcE improved glycopeptide detection by LC-MS. The double-StcE approach maximized glycopeptide recovery from (A) the mucinous tumors, based on HexNAc traces. (B) Strong HexNAc signal was also detected in the non-tumor regions for Colon 1 and Colon 2 prepared with the double-StcE workflow. Regions are numbered according to Figure S5a-f. NL, normalized level.

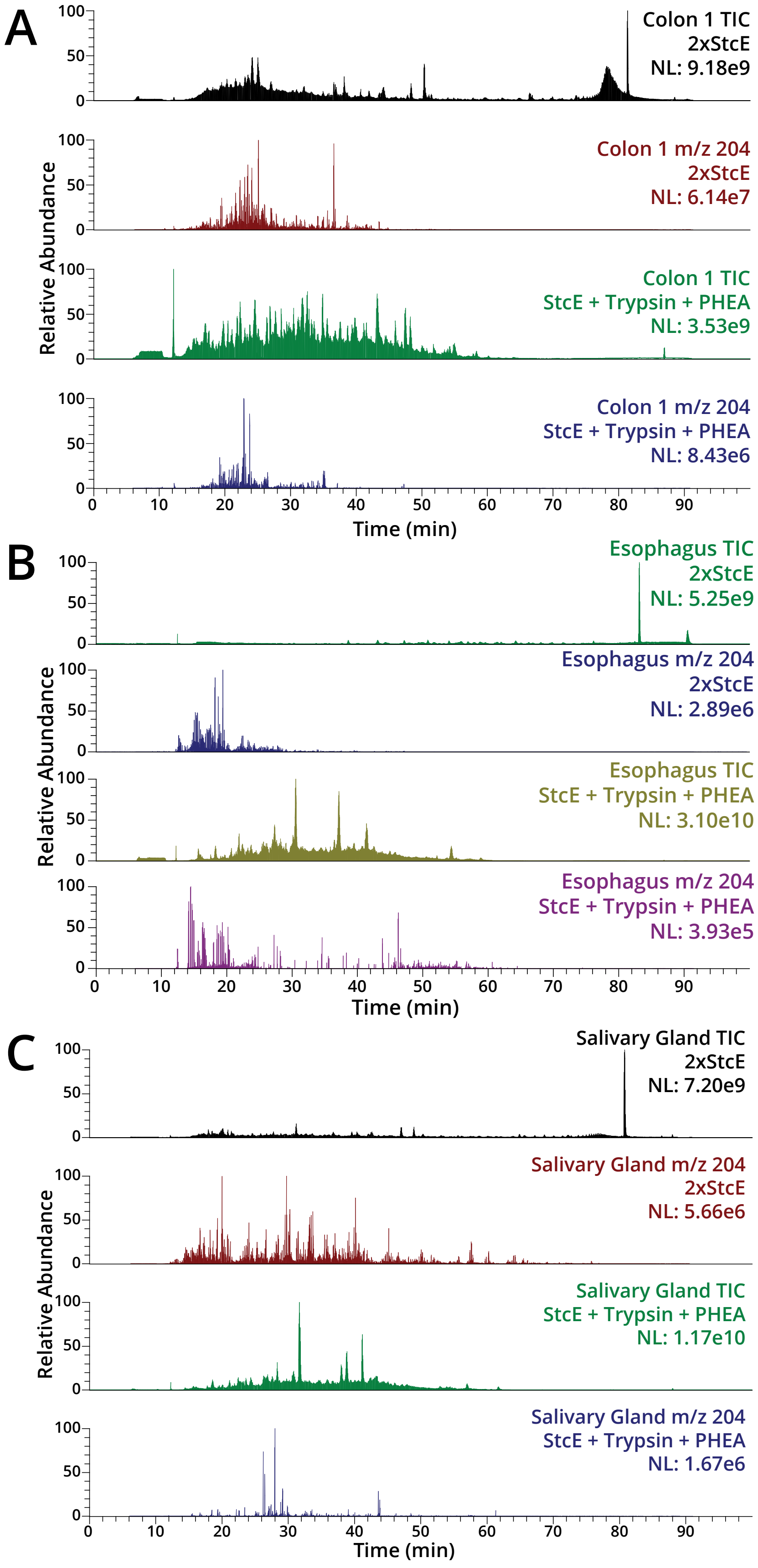
A
Colon 1 TIC
2xStcE
NL: 9.18e9
Colon 1 m/z 204
2xStcE
NL: 6.14e7
Colon 1 TIC
StcE + Trypsin + PHEA
NL: 3.53e9
Colon 1 m/z 204
StcE + Trypsin + PHEA
NL: 8.43e6
Relative Abundance
Time (min)
B
Esophagus TIC
2xStcE
NL: 5.25e9
Esophagus m/z 204
2xStcE
NL: 2.89e6
Esophagus TIC
StcE + Trypsin + PHEA
NL: 3.10e10
Esophagus m/z 204
StcE + Trypsin + PHEA
NL: 3.93e5
Relative Abundance
Time (min)
C
Salivary Gland TIC
2xStcE
NL: 7.20e9
Salivary Gland m/z 204
2xStcE
NL: 5.66e6
Salivary Gland TIC
StcE + Trypsin + PHEA
NL: 1.17e10
Salivary Gland m/z 204
StcE + Trypsin + PHEA
NL: 1.67e6
Relative Abundance
Time (min)

**Fig. S9.** The workflow employing a double-StcE digestion had improved glycopeptide recovery from tumor regions compared to the StcE-trypsin-PHEA method. (A) Total ion chromatograms (TICs) and HexNAc ion traces from Colon 1 tumor region. (B) Total ion chromatograms (TICs) and HexNAc ion traces from esophageal tumor region. (C) Total ion chromatograms (TICs) and HexNAc ion traces from salivary gland tumor region.

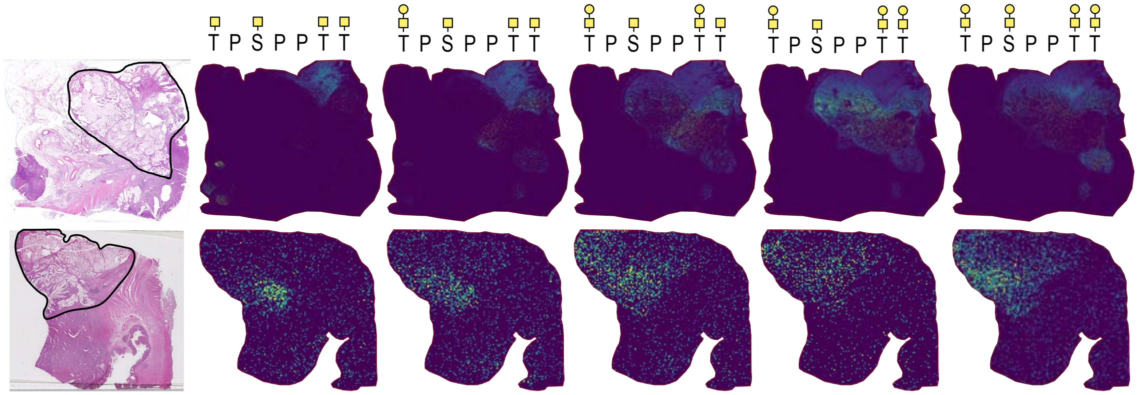


**Fig. S10.** TPSPPTT glycoforms with four glycosites exhibit similar spatial distributions to glycoforms with only three glycosites shown in Figures 2b-c. MALDI-IMS data for PNGaseF- and StcE-treated Colon 1 (top row) was acquired with a timsTOF fleX MALDI-QTOF mass spectrometer (Bruker). Images for PNGaseF- and StcE-treated Colon 2 (bottom row) were acquired with a Solarix dual-source 7T MALDI-FTICR mass spectrometer (Bruker). Each m/z value was manually extracted using SCiLS Lab version 2024b Pro (Bruker) and manually inspected for tumor localization prior to image export. All m/z values shown correspond to the mono-sodiated precursor mass of the associated glycoform. Species were identified using LC-MS data collected for the tumor regions of Colon 1 and 2 prepared with the double-StcE workflow.

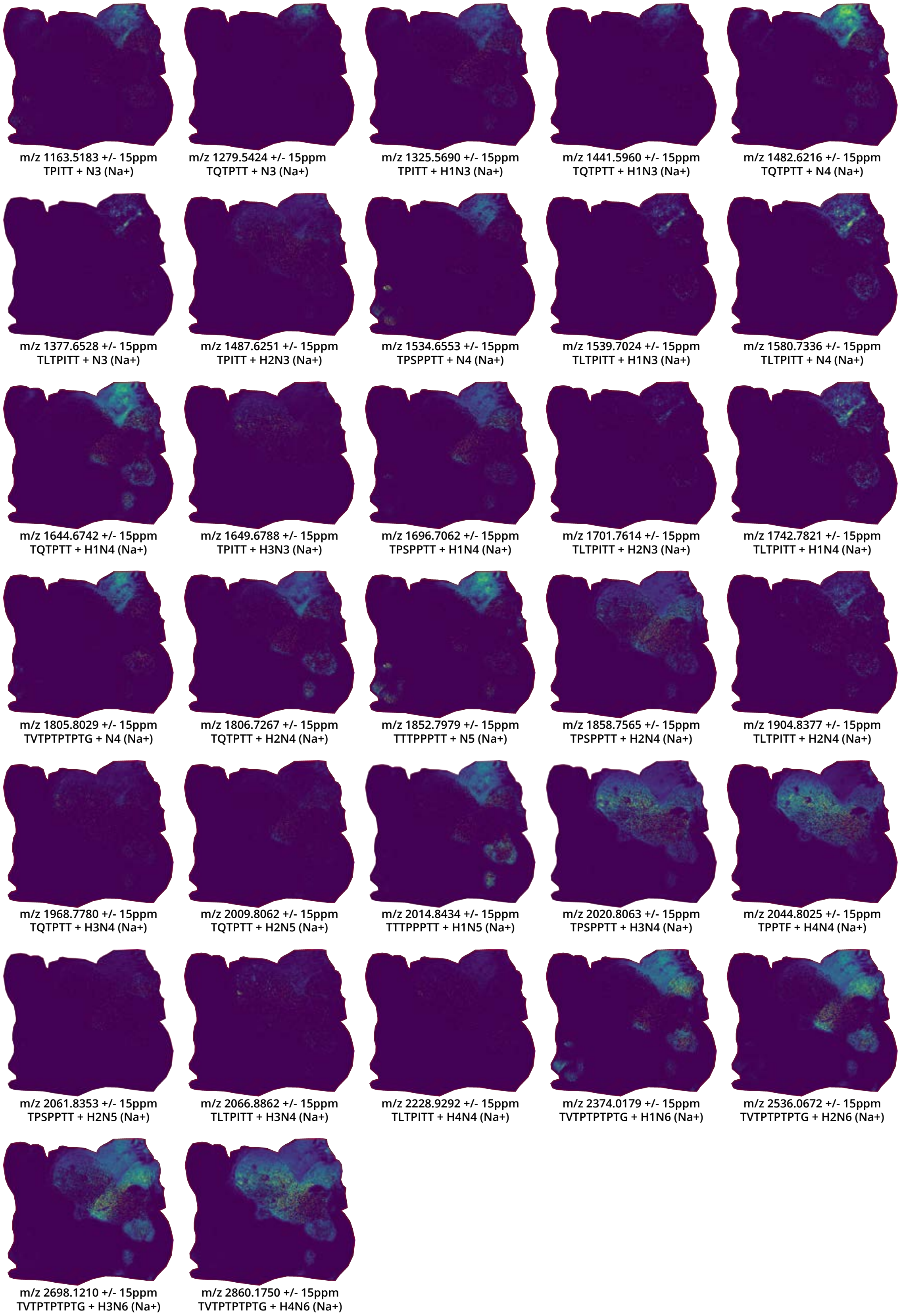
m/z 1163.5183 +/- 15ppm
TPITT + N3 (Na+)
m/z 1279.5424 +/- 15ppm
TQTPTT + N3 (Na+)
m/z 1325.5690 +/- 15ppm
TPITT + H1N3 (Na+)
m/z 1441.5960 +/- 15ppm
TQTPTT + H1N3 (Na+)
m/z 1482.6216 +/- 15ppm
TQTPTT + N4 (Na+)
m/z 1377.6528 +/- 15ppm
TLTPITT + N3 (Na+)
m/z 1487.6251 +/- 15ppm
TPITT + H2N3 (Na+)
m/z 1534.6553 +/- 15ppm
TPSPPTT + N4 (Na+)
m/z 1539.7024 +/- 15ppm
TLTPITT + H1N3 (Na+)
m/z 1580.7336 +/- 15ppm
TLTPITT + N4 (Na+)
m/z 1644.6742 +/- 15ppm
TQTPTT + H1N4 (Na+)
m/z 1649.6788 +/- 15ppm
TPITT + H3N3 (Na+)
m/z 1696.7062 +/- 15ppm
TPSPPTT + H1N4 (Na+)
m/z 1701.7614 +/- 15ppm
TLTPITT + H2N3 (Na+)
m/z 1742.7821 +/- 15ppm
TLTPITT + H1N4 (Na+)
m/z 1805.8029 +/- 15ppm
TVTPTPTPTG + N4 (Na+)
m/z 1806.7267 +/- 15ppm
TQTPTT + H2N4 (Na+)
m/z 1852.7979 +/- 15ppm
TTTPPPTT + N5 (Na+)
m/z 1858.7565 +/- 15ppm
TPSPPTT + H2N4 (Na+)
m/z 1904.8377 +/- 15ppm
TLTPITT + H2N4 (Na+)
m/z 1968.7780 +/- 15ppm
TQTPTT + H3N4 (Na+)
m/z 2009.8062 +/- 15ppm
TQTPTT + H2N5 (Na+)
m/z 2014.8434 +/- 15ppm
TTTPPPTT + H1N5 (Na+)
m/z 2020.8063 +/- 15ppm
TPSPPTT + H3N4 (Na+)
m/z 2044.8025 +/- 15ppm
TPPTF + H4N4 (Na+)
m/z 2061.8353 +/- 15ppm
TPSPPTT + H2N5 (Na+)
m/z 2066.8862 +/- 15ppm
TLTPITT + H3N4 (Na+)
m/z 2228.9292 +/- 15ppm
TLTPITT + H4N4 (Na+)
m/z 2374.0179 +/- 15ppm
TVTPTPTPTG + H1N6 (Na+)
m/z 2536.0672 +/- 15ppm
TVTPTPTPTG + H2N6 (Na+)
m/z 2698.1210 +/- 15ppm
TVTPTPTPTG + H3N6 (Na+)
m/z 2860.1750 +/- 15ppm
TVTPTPTPTG + H4N6 (Na+)

**Fig. S11.** Additional glycopeptides detected by MALDI-MSI in the tumor region of PNGaseF- and StcE-treated Colon 1 that were subsequently identified with LC-MS data. Images were acquired with a timsTOF fleX MALDI-QTOF (Bruker). Each m/z value was manually extracted using SCiLS Lab version 2024b Pro (Bruker) and manually inspected for tumor localization prior to image export. All m/z values shown correspond to the mono-sodiated precursor mass of the associated glycoform. Species were identified using LC-MS data collected for the tumor region of Colon 1 prepared with the double-StcE workflow.

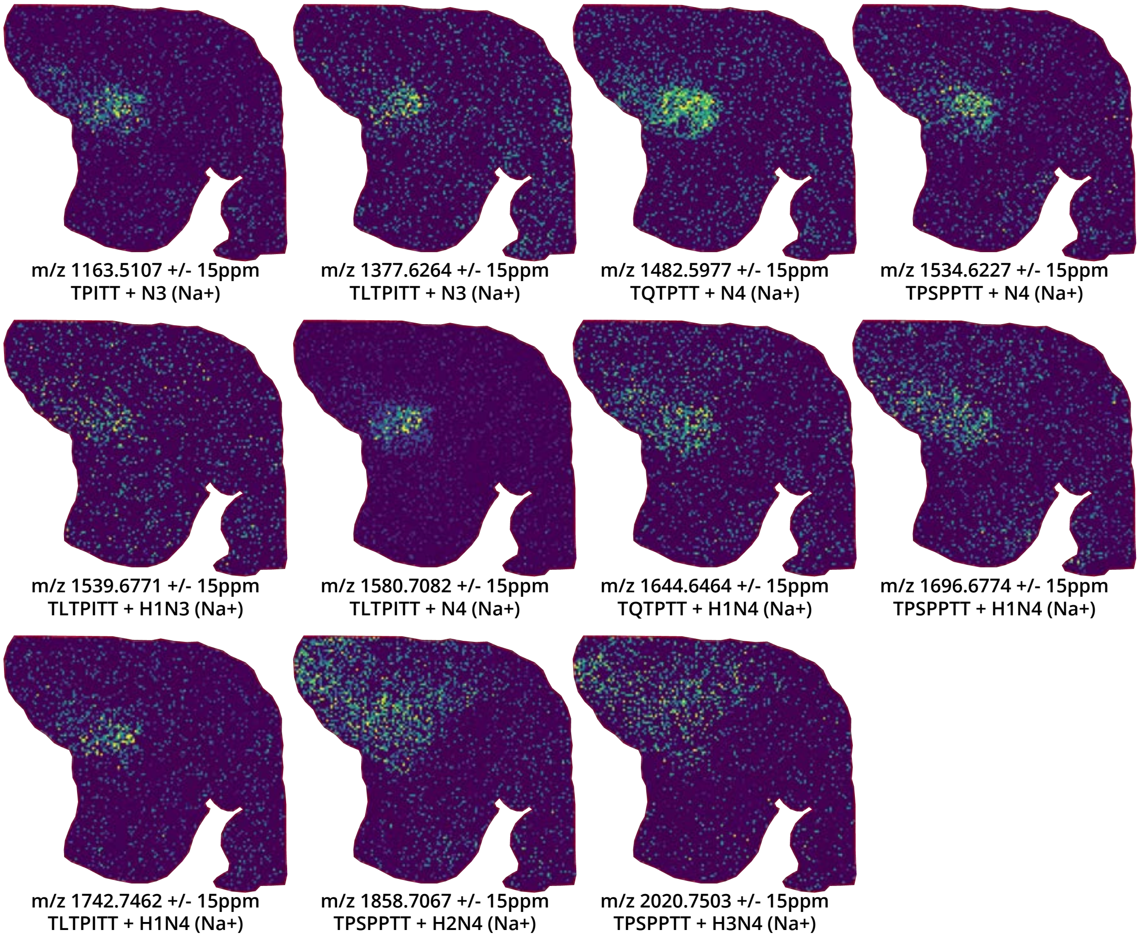


**Fig. S12.** Additional glycopeptides detected by MALDI-MSI in the tumor region of PNGaseF- and StcE-treated Colon 2 that were subsequently identified by LC-MS. Images were acquired with a Solarix dual-source 7T MALDI-FTICR mass spectrometer (Bruker). Each m/z value was manually extracted using SCiLS Lab version 2024b Pro (Bruker) and manually inspected for tumor localization prior to image export. The m/z values shown correspond to mono-sodiated precursor masses for the associated glycoform. Species were identified using LC-MS data collected for the tumor region of Colon 2 prepared with the double-StcE workflow.

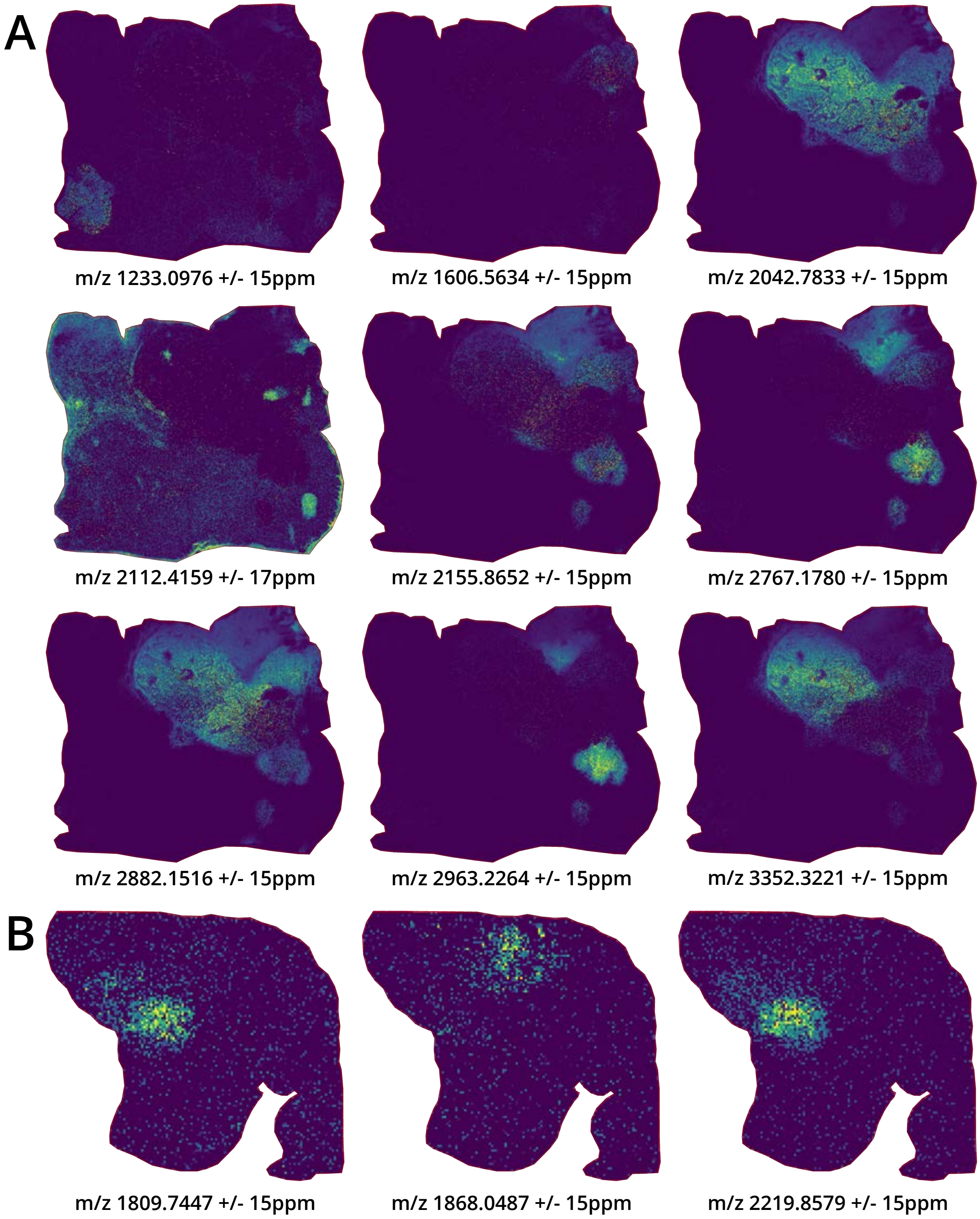


**Fig. S13.** Tumor-associated ions detected in PNGaseF- and StcE-treated colorectal tumors by MALDI-MSI that were not identified with LC-MS data. Each m/z value was manually extracted in SCiLS Lab version 2024b Pro (Bruker) and assessed for tumor localization prior to image export. (A) Nine unidentifiable species were detected at a strong intensity in Colon 1. Images were acquired with a timsTOF fleX MALDI-QTOF mass spectrometer. (B) Three unidentifiable species detected at a strong intensity in Colon 2. Images were acquired with a Solarix dual-source 7T MALDI-FTICR mass spectrometer (Bruker).

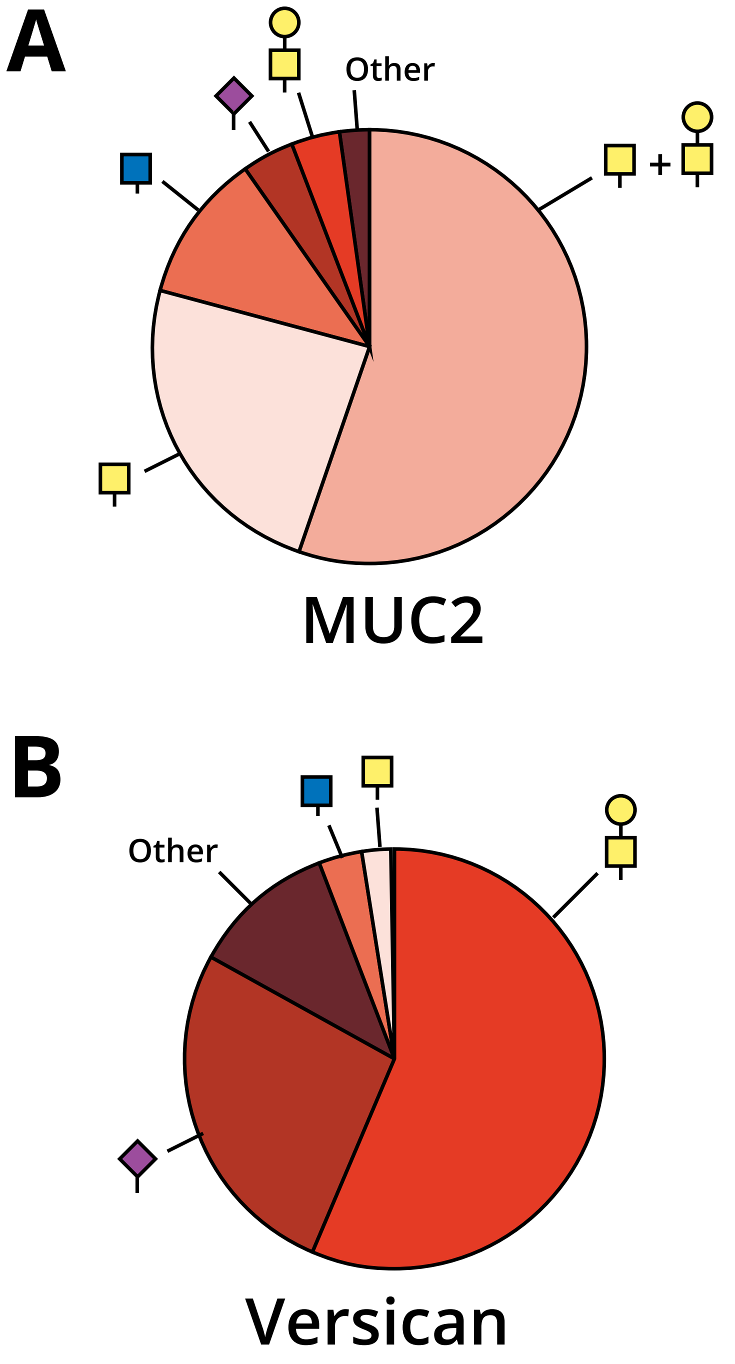


**Fig. S14.** Pie charts showing distribution of identified glycoforms. (A) MUC2 glycopeptides (N=553) can be categorized into Tn+T (306, 55.3%), Tn-only (133, 24.1%), GlcNAc-containing (61, 11.0%), sialylated (22, 4.0%), T-only (20, 3.6%), and other (11, 2.0%) species. (B) Versican glycopeptides (N=251) can be categorized into T-only (142, 56.6%), sialylated (67, 26.7%), other (28, 11.2%), GlcNAc-containing (8, 3.2%), and Tn-containing (6, 2.4%) species.

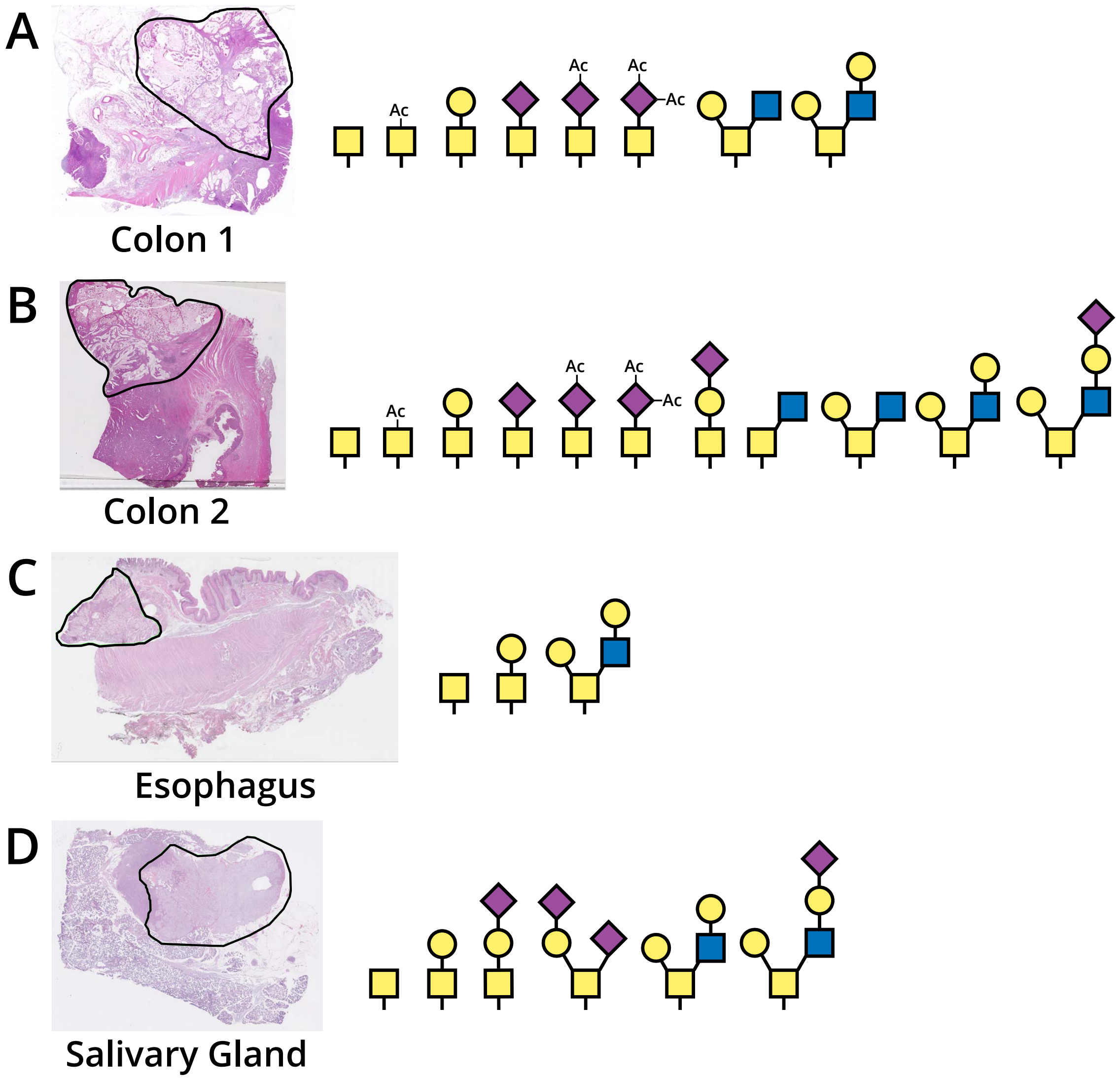


**Fig. S15.** Glycan structures confidently localized in each tumor region (outlined in black) via LC-MS/MS. (A) From left to right: N1 (Tn), AcN1, H1N1 (T), N1A1 (sialyl-Tn), N1AcA1, N1diAcA1, H1N2, and H2N2 could be localized to specific sites on Colon 1 glycopeptides. (B) From left to right: N1 (Tn), AcN1, H1N1 (T), N1A1 (sialyl-Tn), N1AcA1, N1diAcA1, H1N1A1 (sialyl T), N2, H1N2, H2N2, and H2N2A1 could be localized to specific sites on Colon 2 glycopeptides. (C) From left to right: N1 (Tn), H1N1 (T), and H2N2 could be localized to specific sites on Esophagus glycopeptides. (D) From left to right: N1 (Tn), H1N1 (T), H1N1A1 (sialyl T), H1N1A2 (di-sialyl T), H2N2, and H2N2A1 could be localized to specific sites on Salivary Gland glycopeptides.

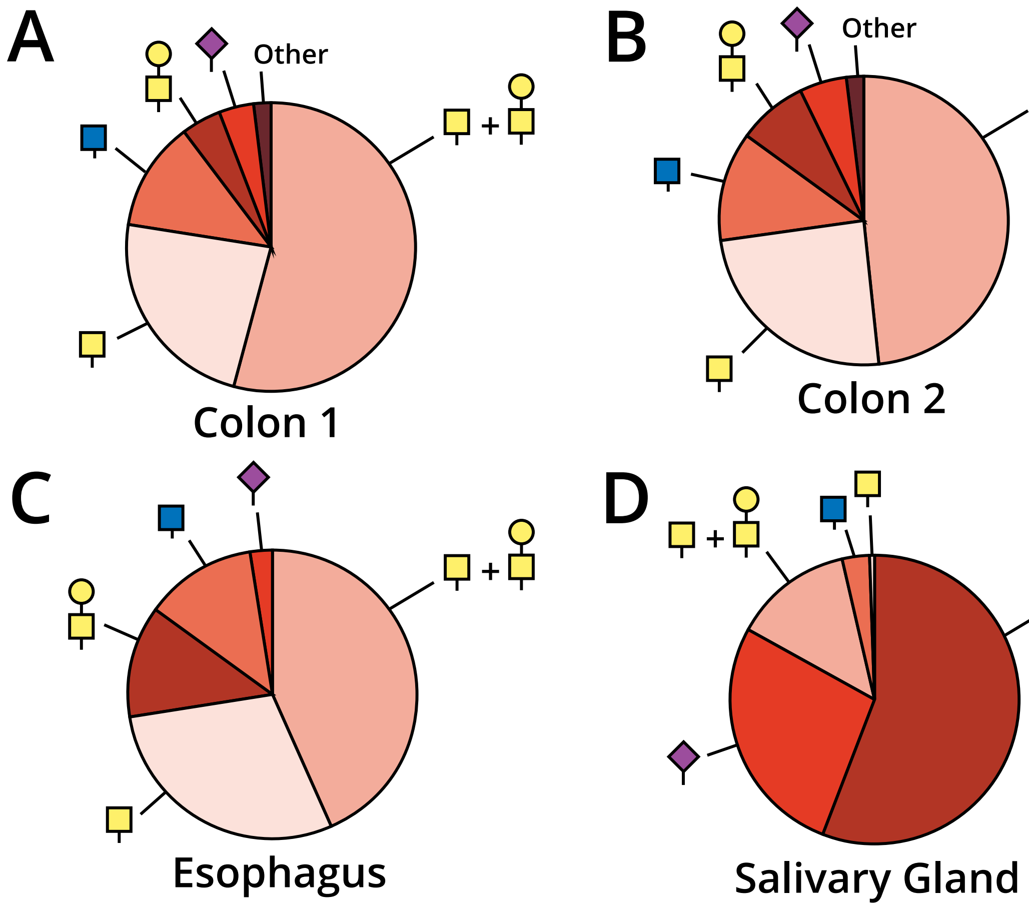


**Fig. S16.** Pie charts showing distribution of glycoforms identified in each tumor region. (A) Colon 1 glycopeptides (N=565) can be categorized into Tn+T (306, 54.2%), Tn-only (132, 23.4%), GlcNAc-containing (69, 12.2%), T-only (25, 4.4%), sialylated (22, 3.9%), and other (11, 1.9%) species. (B) Colon 2 glycopeptides (N=575) can be categorized into Tn+T (279, 48.5%), Tn-only (140, 24.4%), GlcNAc-containing (70, 12.2%), T-only (45, 7.8%), sialylated (30, 5.2%), and other (11, 1.9%) species. (C) Esophagus glycopeptides (N=216) can be categorized into Tn+T (94, 43.5%), Tn-only (63, 29.2%), GlcNAc-containing (27, 12.5%), T-only (27, 12.5%), and sialylated (5, 2.3%) species. (D) Salivary gland glycopeptides (N=254) can be categorized into T-only (142, 55.9%), sialylated (69, 27.2%), Tn+T (34, 13.4%), GlcNAc-containing (8, 3.1%), and Tn-only (1, 0.4%) species.

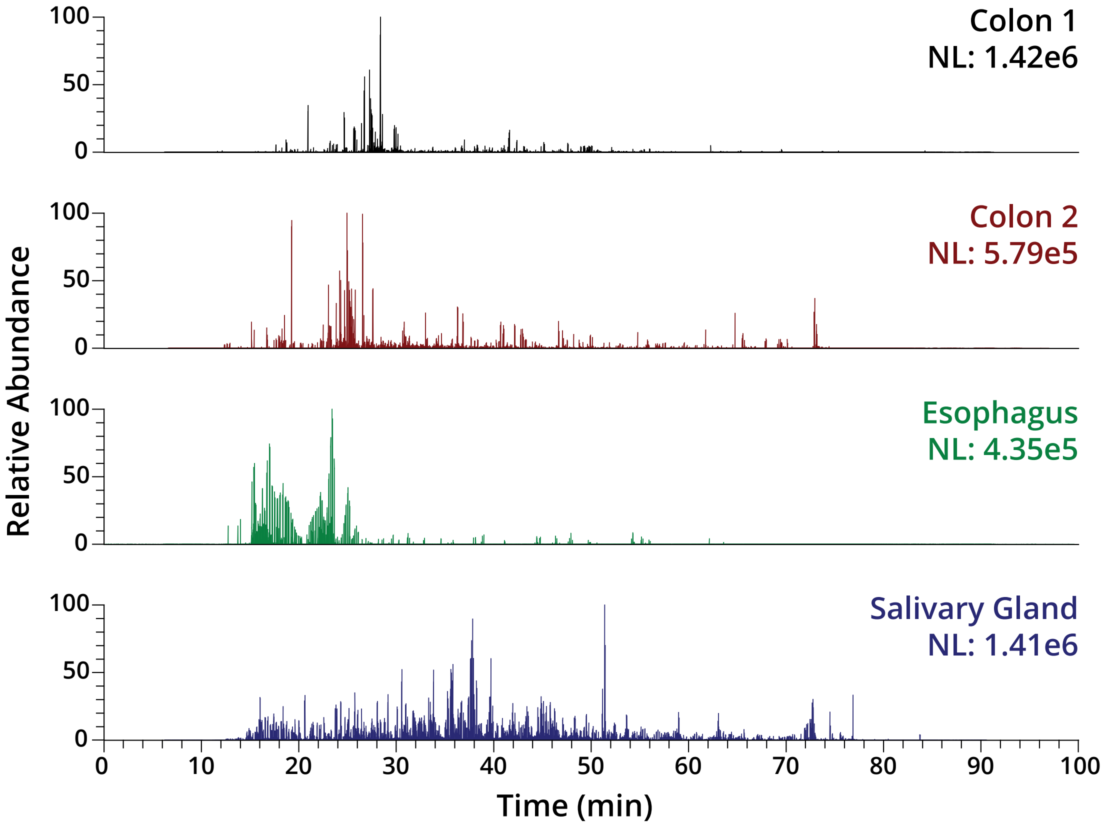


**Fig. S17.** Combined traces of Neu5Ac fingerprint ions at *m/z* 292.1027 and 274.0921 in the tumor region of Colon 1 (black), Colon 2 (red), esophagus (green), and salivary gland (blue). All samples were prepared with the double-StcE workflow. NL, normalized level.

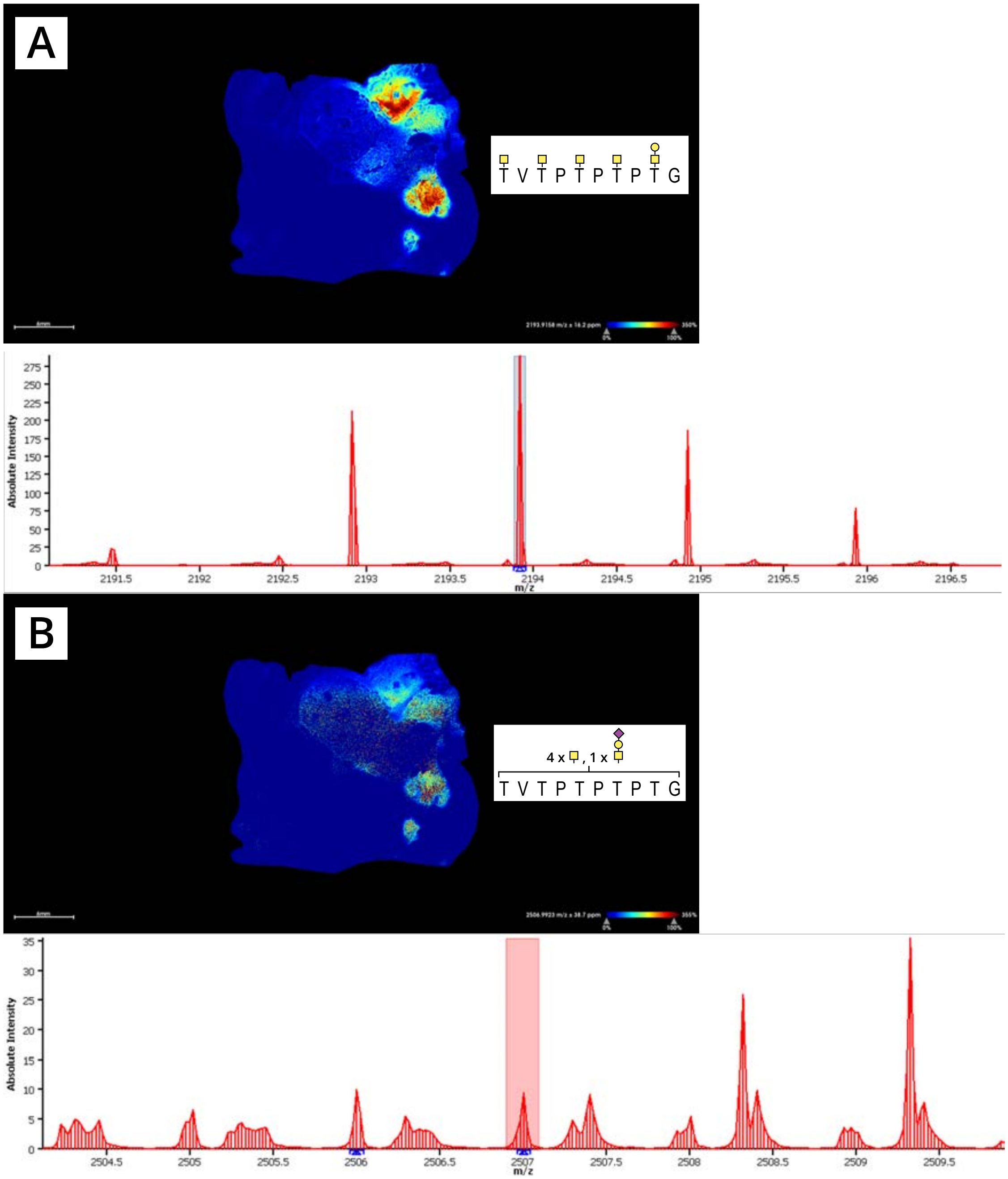


**Fig. S18.** Sialylated MUC2 glycoforms were less intense than their non-sialylated counterparts in Colon 1 MALDI-MSI experiments. (A) Heat map of glycopeptide with backbone sequence TVTPTPTPTG from the second PTS domain of MUC2 decorated with one T and four Tn antigen structures. Note that this intensity map corresponds to the isotopic peak with a single C13 atom (bottom row), which is more intense than the monoisotopic precursor mass. (B) Heat map of glycopeptide with backbone sequence TVTPTPTPTG from the second PTS domain of MUC2 decorated with one ST and four Tn antigen structures. For consistency with Panel A, the C13 peak of this species was also used to generate the intensity map, although the monoisotopic precursor mass is at roughly the same intensity (bottom row). MALDI-MSI data was acquired with a timsTOF fleX MALDI-QTOF mass spectrometer (Bruker). Ion images were manually extracted in SCiLS Lab version 2024b (Bruker).

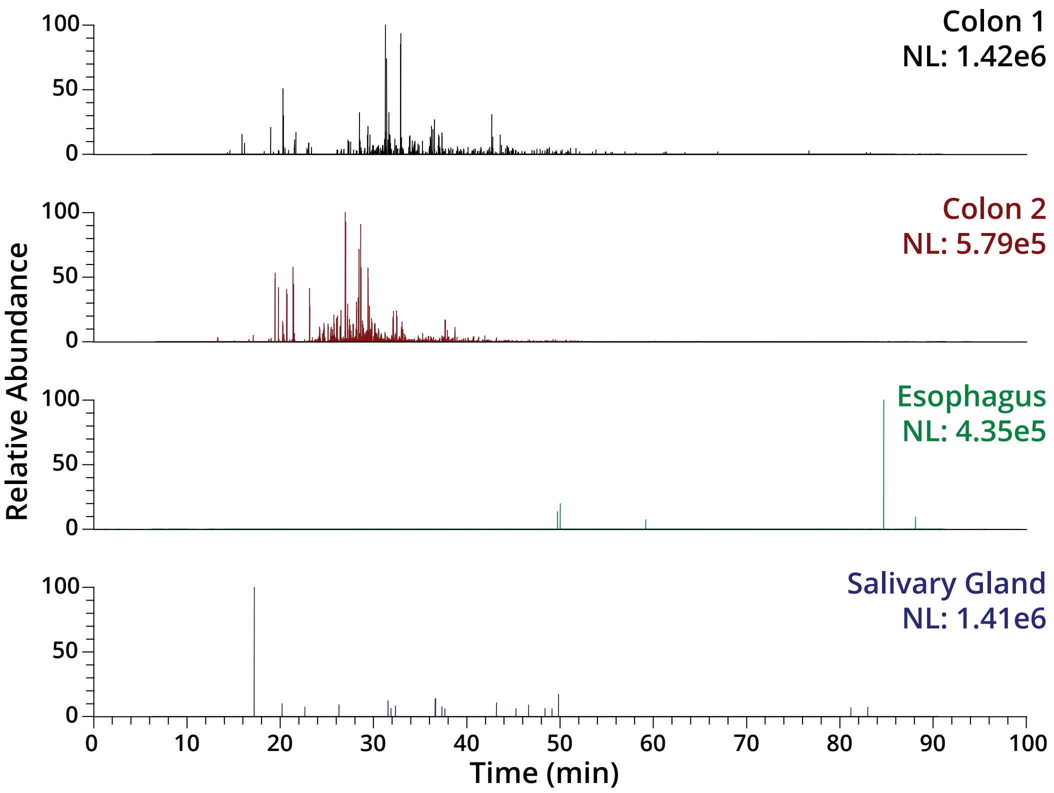


**Fig. S19.** Combined traces of AcNeu5Ac fingerprint ions at *m/z* 334.1132 and 316.1026 for the tumor regions of Colon 1 (black), Colon 2 (red), esophagus (green), and salivary gland (blue). All samples shown were prepared with the double-StcE workflow. NL, normalized level.

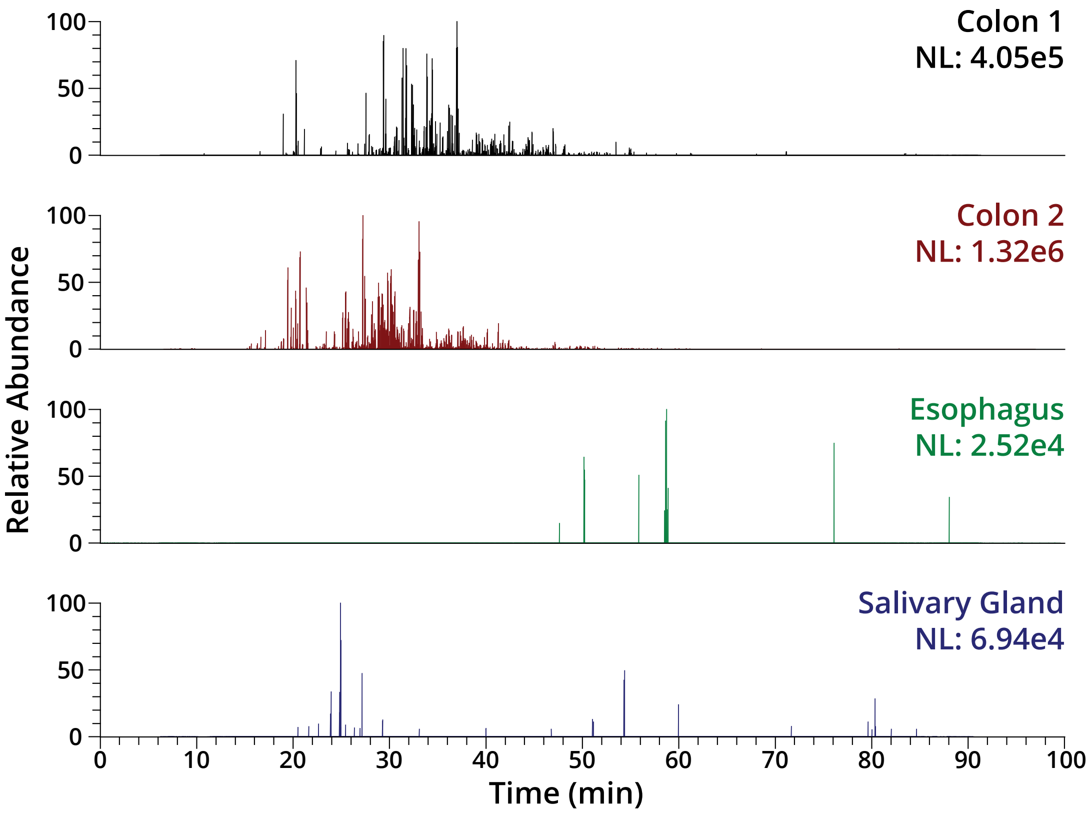


**Fig. S20.** Combined traces of $Ac_2Neu5Ac$ fingerprint ions at *m/z* 376.1237 and 358.1131 for the tumor regions of Colon 1 (black), Colon 2 (red), esophagus (green), and salivary gland (blue). All samples shown were prepared with the double-StcE workflow. NL, normalized level.

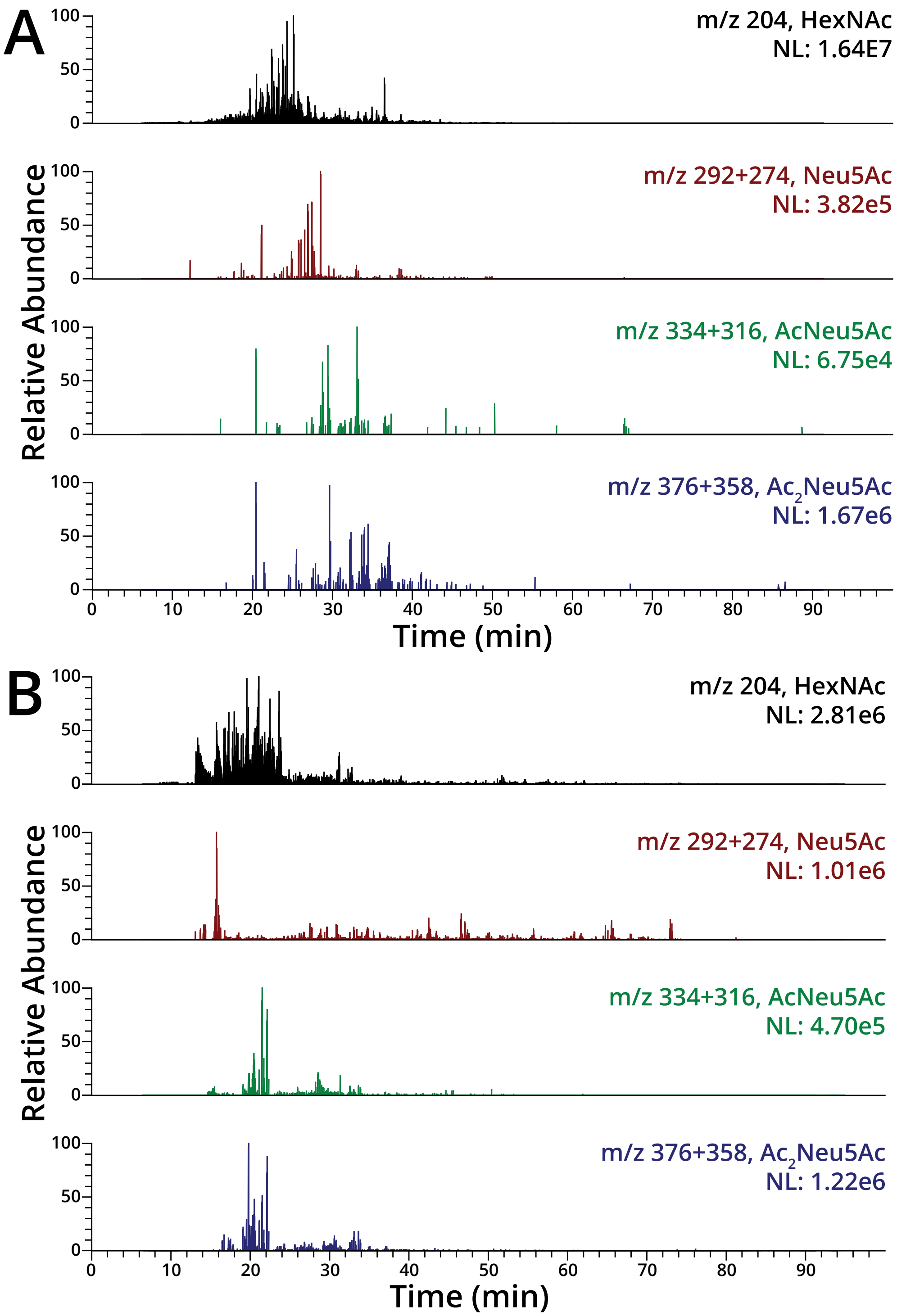


**Fig. S21.** Traces of AcNeu5Ac and $Ac_2Neu5Ac$ fingerprint ions in non-tumor regions. (A) Marker ion traces for HexNAc (black), Neu5Ac (red), AcNeu5Ac (green), and $Ac_2Neu5Ac$ (blue) for the non-tumor region of Colon 1 prepared with the double-StcE workflow. (B) Marker ion traces for

HexNAc (black), Neu5Ac (red), AcNeu5Ac (green), and $Ac_2$Neu5Ac (blue) for the non-tumor region of Colon 2 prepared with the double-StcE workflow. NL, normalized level.

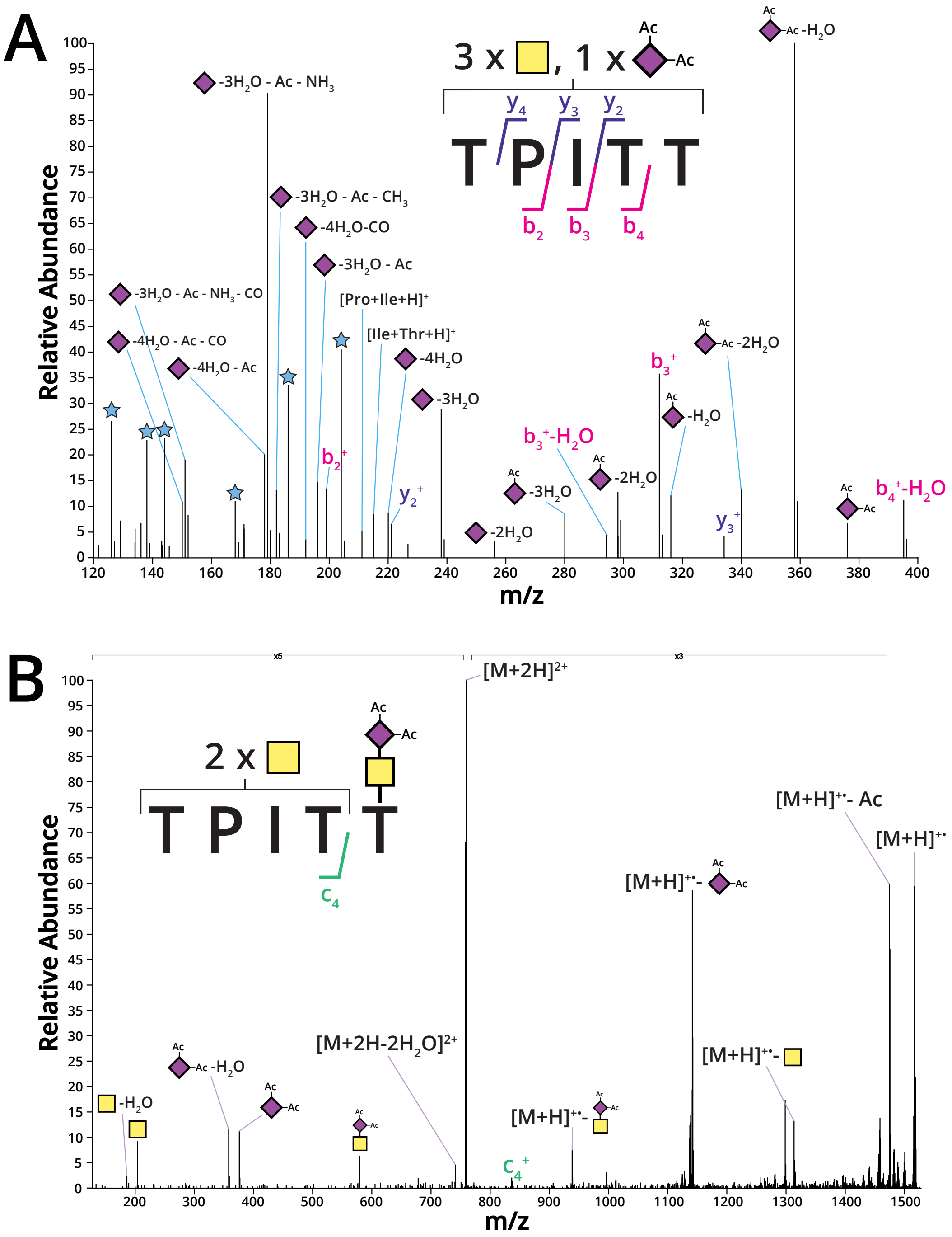


**Fig. S22.** Additional MS2 evidence for Ac$_2$Neu5Ac-containing glycopeptide. These spectra are from the non-tumor region data for Colon 2 prepared with the double-StcE workflow and are representative of other spectra for Ac$_2$Neu5Ac species. (A) Low-mass region (*m/z* 120-400) of the higher-energy collision dissociation (HCD) spectrum shown in Figure 4A for the MUC2 glycopeptide with sequence TPITT, demonstrating the rich fragmentation of the Ac$_2$Neu5Ac moiety. Blue stars represent HexNAc fingerprint ions. (B) Electron-transfer dissociation (ETD) spectrum for the same precursor ion.

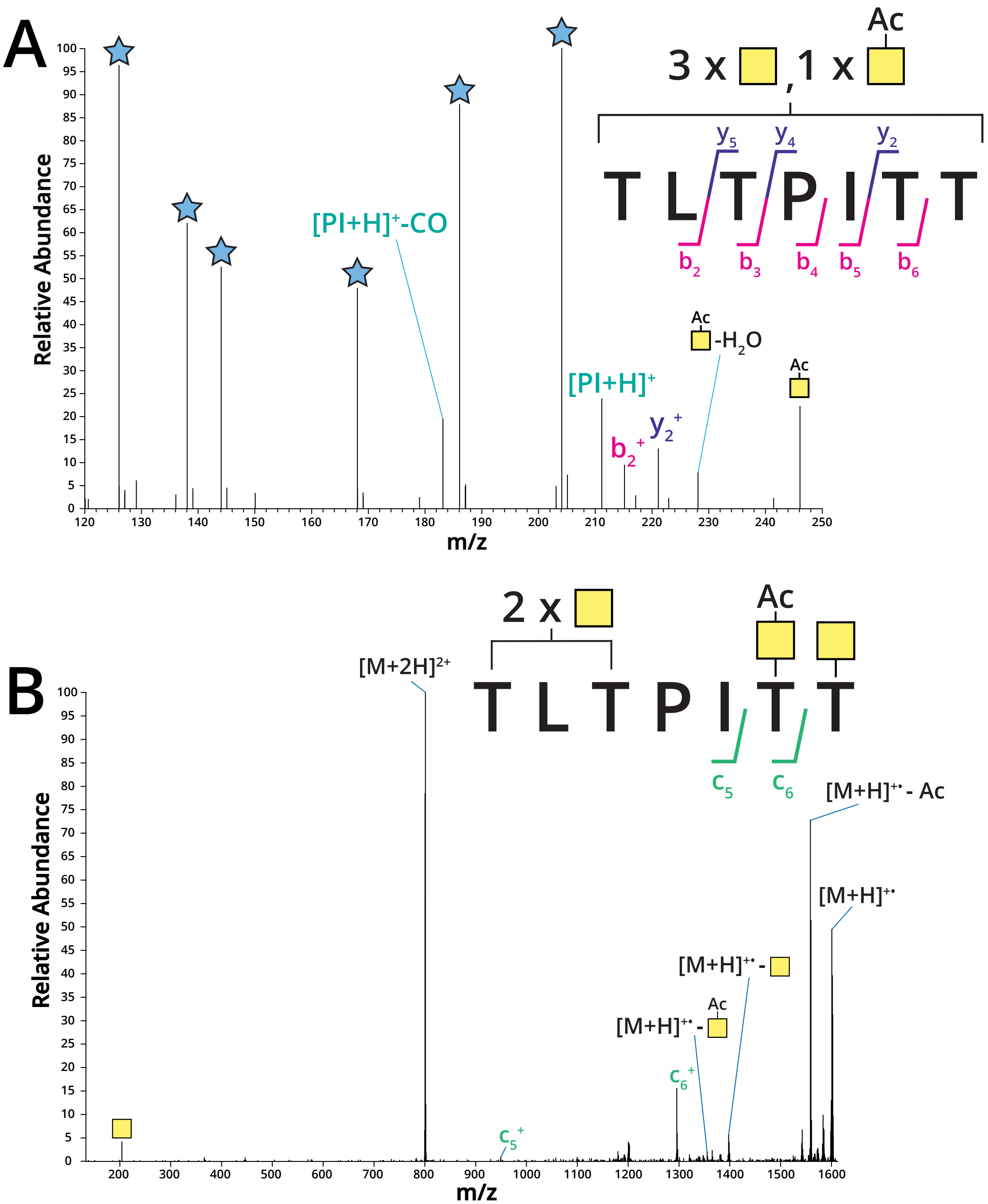


**Fig. S23.** Additional MS2 evidence for glycopeptide decorated with acetylated GalNAc. (A) Low-mass region of HCD spectrum shown in Figure 4B for the MUC2 glycopeptide with sequence TLTPITT, showing fragmentation of a putative AcGalNAc-containing glycopeptide. Marker ions for the intact AcGalNAc structure as well as its water loss were detected. (B) ETD spectrum for the same precursor. Spectra are from tumor region LC-MS data for Colon 2 prepared with the double-StcE workflow.

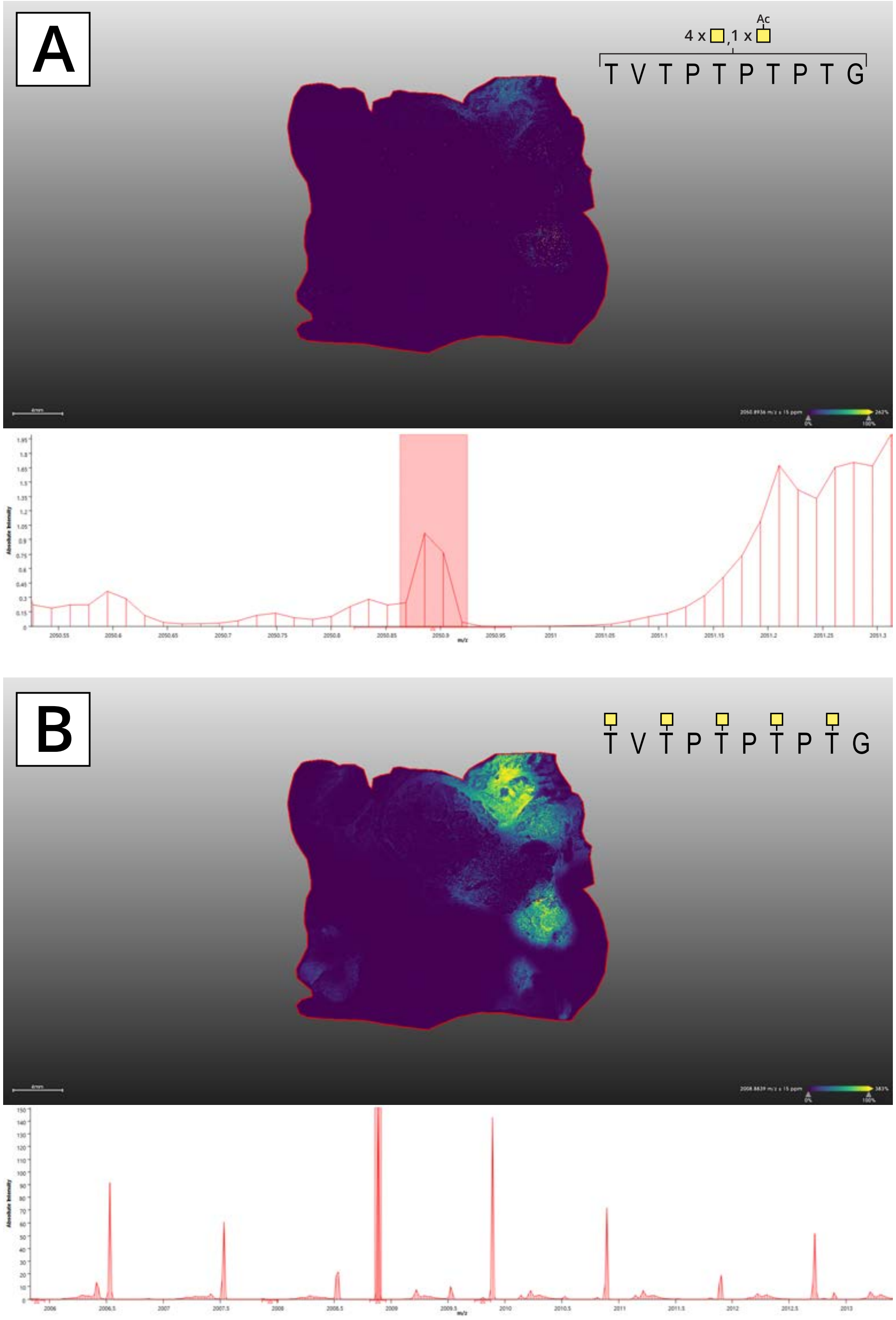


**Fig. S24.** A MUC2 glycoform with AcGalNAc overlaps with Tn-only glycoform expression in MALDI-MSI of Colon 1. This tissue section was treated with PNGaseF and StcE prior to data acquisition with a timsTOF fleX MALDI-QTOF mass spectrometer (Bruker). Precursor ions detected in the tumor region of double-StcE-treated Colon 1 with LC-MS were converted to mono-sodiated m/z values, which were manually extracted in MALDI-IMS data using SCiLS Lab version 2024b Pro (Bruker). (A) Heat map for the mass corresponding to the glycopeptide TVTPTPTPTG containing four Tn antigens and one AcGalNAc structure, which had a very low overall intensity (bottom row). This spatial distribution overlapped with the highest expression of the Tn-only glycoform shown in (B), which had a much higher overall abundance (bottom row).

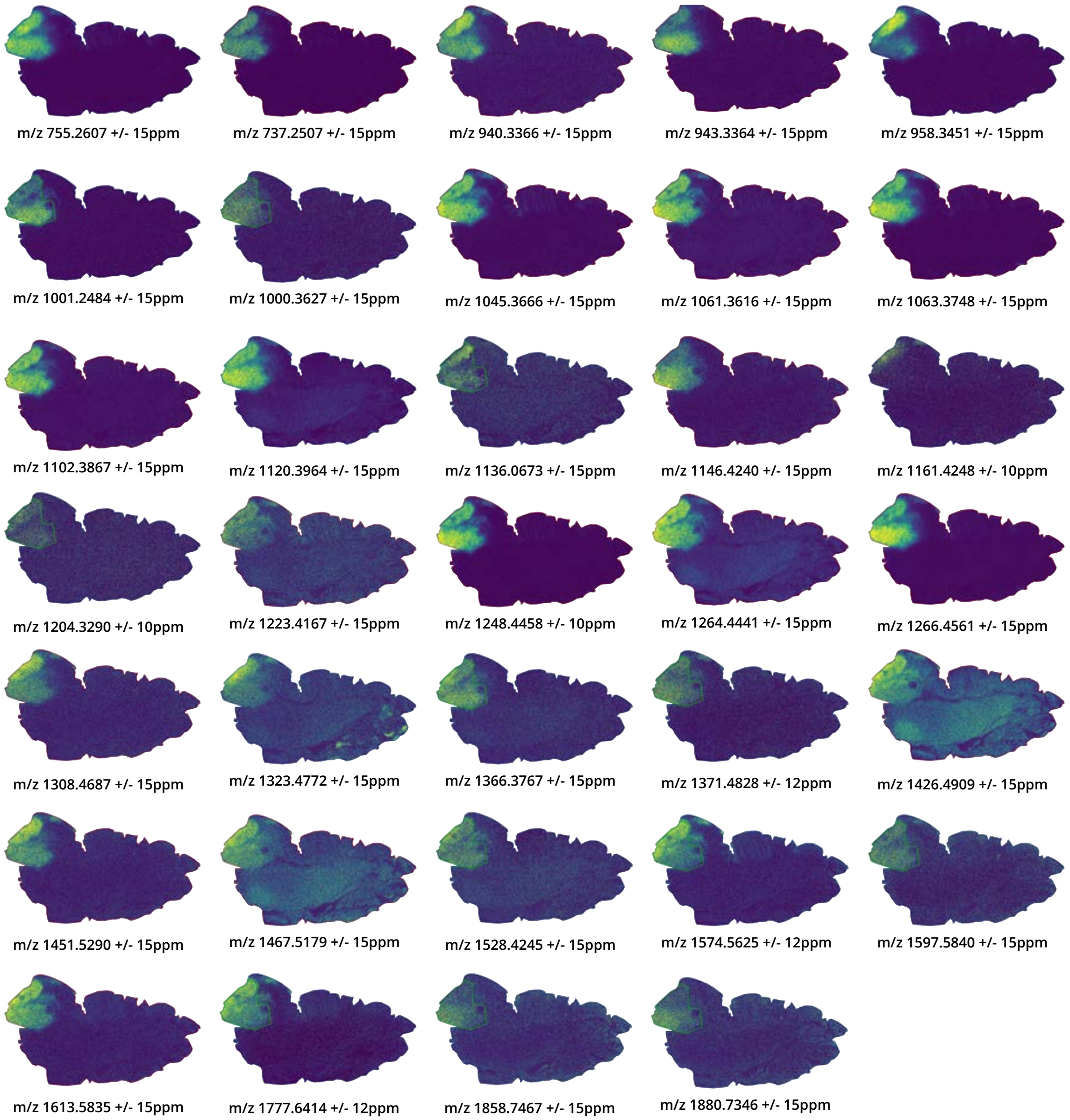


**Fig. S25.** Tumor-associated ions detected by MALDI-MSI of the PNGaseF- and StcE-treated esophageal tissue. Raw data was acquired with a timsTOF fleX MALDI-QTOF mass spectrometer (Bruker). Each m/z value was manually extracted in SCiLS Lab version 2024b Pro (Bruker) and inspected for tumor localization prior to image export. The depicted species could not be identified with corresponding LC-MS data.

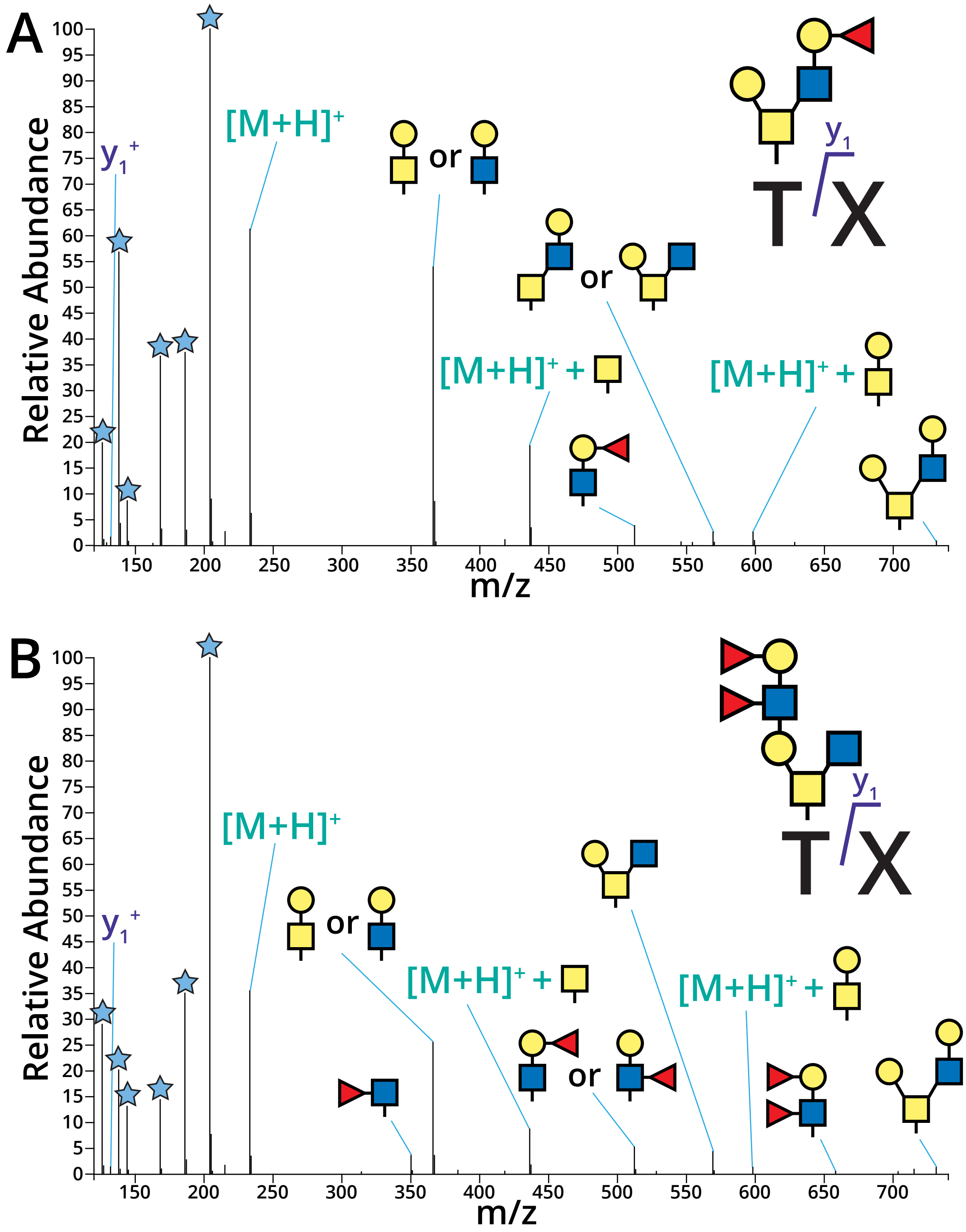


**Fig. S26.** O-Glycosylated dipeptides were detected by LC-MS in the tumor region of the esophageal tissue prepared with the double-StcE workflow. (A) HCD spectrum for the ion at a retention time of 18.22 minutes with m/z 555.2397, corresponding to the +2 precursor of dipeptide TL or TI decorated with a mono-fucosylated core 2 O-glycan. (B) HCD spectrum for the ion at a retention time of 19.41 minutes with m/z 729.8077, corresponding to the +2 precursor of dipeptide TL or TI decorated with a core 2 O-glycan likely extended with a $Le^b$ or $Le^y$ motif. Blue stars in low-mass region correspond to HexNAc fingerprint ions.

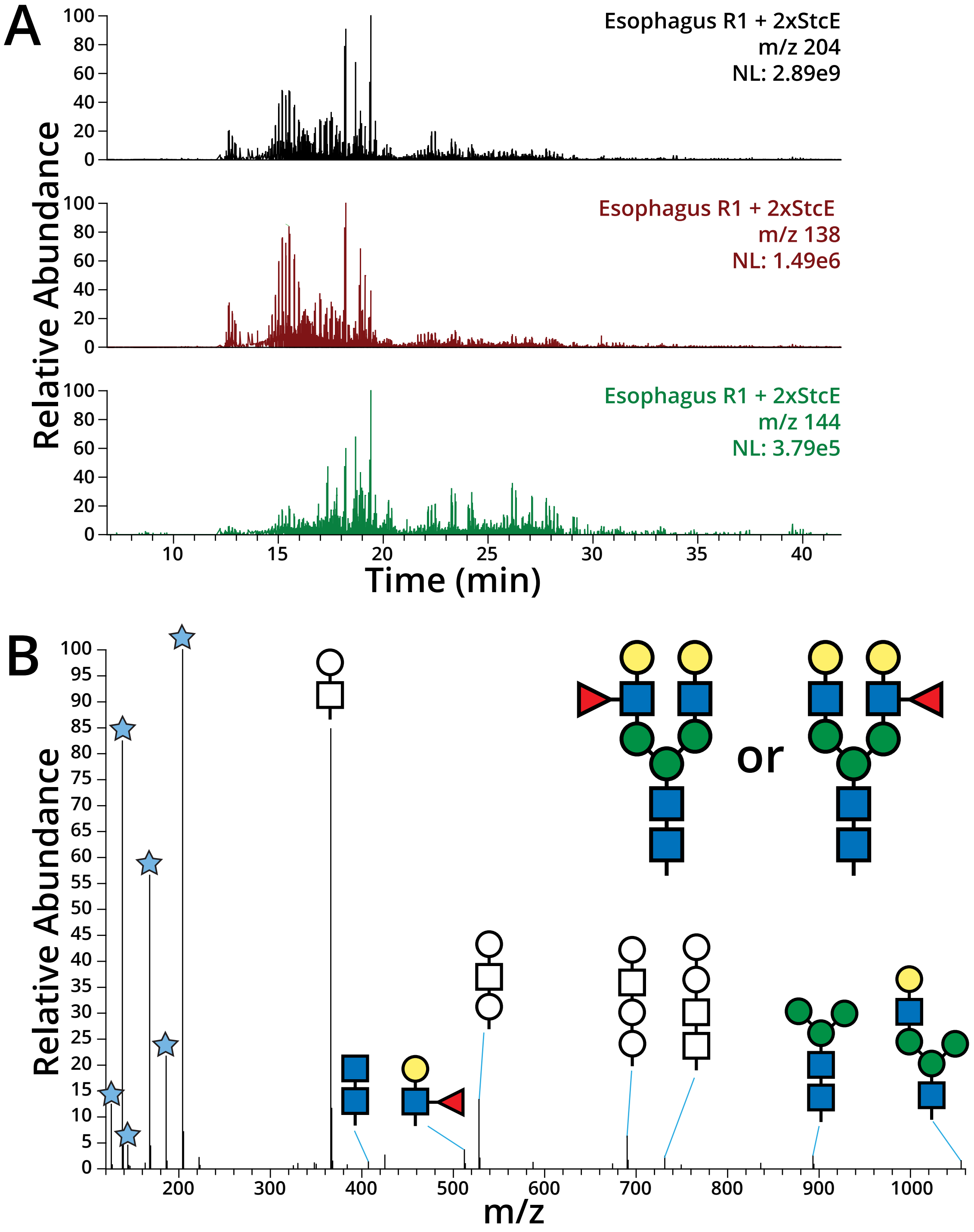


**Fig. S27.** Free N-glycans were relatively abundant in the LC-MS data for the double-StcE-treated esophageal tumor region. (A) Traces of the HexNAc marker ions at m/z 204.0867 (black), 138.055 (red), and 144.0655 (green). (B) HCD spectrum for the ion at a retention time of 15.19 minutes and with m/z 894.3327, corresponding to the +2 precursor for the N-glycan with composition H5N4F1. Uncolored shapes indicate glycan fragments that cannot be attributed to a specific portion of the N-glycan structure. Blue stars in the low-mass region correspond to HexNAc fingerprint ions.

**Table S1.** Tumor-associated proteins identified in unmodified peptide data. Proteome results were filtered for proteins found in all tumors, but not in any samples collected from non-tumor regions. C1, Colon 1; C2, Colon 2; E, Esophagus; SG, Salivary Gland. "-T" indicates tumor region, and "-N" denotes non-tumor region.

| UniProt ID | Description | C1-T | C1-N | C2-T | C2-N | E-T | E-N | SG-T | SG-N |
|---|---|---|---|---|---|---|---|---|---|
| Q14767 | Latent-transforming growth factor beta-binding protein 2 | + | - | + | - | + | - | + | - |
| P08621 | U1 small nuclear ribonucleoprotein 70 kDa | + | - | + | - | + | - | + | - |
| Q8N163 | Cell cycle and apoptosis regulator protein 2 | + | - | + | - | + | - | + | - |
| P04439 | HLA class I histocompatibility antigen, A alpha chain | + | - | + | - | + | - | + | - |
| Q13263 | Transcription intermediary factor 1-beta | + | - | + | - | + | - | + | - |
| Q9ULA0 | Aspartyl aminopeptidase | + | - | + | - | + | - | + | - |
| Q13510 | Acid ceramidase | + | - | + | - | + | - | + | - |
| P82979 | SAP domain-containing ribonucleoprotein | + | - | + | - | + | - | + | - |

**Table S2.** Extracellular proteases and peptidases identified in unmodified peptide data. C1, Colon 1; C2, Colon 2; E, Esophagus; SG, Salivary Gland. "-T" indicates tumor region, and "-N" denotes non-tumor region.

| UniProt ID | Description | C1-T | C1-N | C2-T | C2-N | E-T | E-N | SG-T | SG-N |
|---|---|---|---|---|---|---|---|---|---|
| P61626 | Lysozyme C | + | + | + | + | + | + | + | + |
| P07339 | Cathepsin D | + | + | + | + | + | - | + | + |
| P08311 | Cathepsin G | + | + | + | + | + | + | + | - |
| P15088 | Mast cell carboxypeptidase A | - | + | - | + | + | + | + | - |
| P14780 | Matrix metalloproteinase-9 | - | + | + | + | + | - | + | - |
| Q9H4A4 | Aminopeptidase B | - | - | + | + | - | - | + | + |
| P15144 | Aminopeptidase N | - | - | - | - | - | - | + | - |
| P20231 | Tryptase beta-2 | + | + | + | + | + | + | - | - |
| P16444 | Dipeptidase 1 | + | - | + | - | - | - | - | - |
| P17655 | Calpain-2 catalytic subunit | + | - | + | + | - | - | - | - |
| P04632 | Calpain small subunit 1 | + | + | + | + | - | + | - | + |
| P25774 | Cathepsin S | + | - | - | - | - | - | - | - |
| P12955 | Xaa-Pro dipeptidase | + | - | - | - | - | - | - | - |
| P23946 | Chymase | - | + | - | + | + | + | - | - |
| P08253 | 72 kDa type IV collagenase | - | + | - | - | - | - | | |
| P24158 | Myeloblastin | - | - | + | + | - | - | - | - |
| Q96IY4 | Carboxypeptidase B2 | - | - | + | - | - | - | - | - |
| P08217 | Chymotrypsin-like elastase family member 2A | - | - | + | - | - | - | - | - |
| Q92542 | Nicastrin | - | - | + | - | - | - | - | - |
| Q9H3G5 | Probable serine carboxypeptidase CPVL | - | - | - | - | + | - | - | - |
| Q9UHL4 | Dipeptidyl peptidase 2 | - | - | - | - | + | - | - | - |
| P27487 | Dipeptidyl peptidase 4 | - | - | - | - | - | - | - | + |
| O75976 | Carboxypeptidase D | - | - | - | - | - | - | - | + |

**Dataset S1 (separate file).** Proteins identified in unmodified peptide searches.

**Dataset S2 (separate file).** Glycopeptides identified in tumor region of each tissue.
